# Diet-regulated production of PDGFcc by macrophages controls energy storage

**DOI:** 10.1101/2020.06.15.152397

**Authors:** Nehemiah Cox, Lucile Crozet, Inge R. Holtman, Pierre-Louis Loyher, Tomi Lazarov, Elvira Mass, E. Richard Stanley, Christopher K. Glass, Frederic Geissmann

## Abstract

Macrophages control inflammation in obese animals, and may also directly or indirectly regulate energy storage. In a genetic screen we identify a PDGF-family growth factor, *Pvf3*, produced by macrophages and required for lipid storage in *Drosophila* larvae’s fat body cells. We next demonstrate using genetic and pharmacological approaches that *Pvf3* ortholog PDGFcc, produced by *Ccr2*-independent embryo-derived tissue macrophages, is also required for storage in mammalian white adipose tissue. PDGFcc production by resident macrophages is regulated by diet, acts on white adipocytes in a paracrine manner, and controls adipocyte hypertrophy in high-fat diet fed and genetically hyperphagic mice. Upon PDGFcc blockade, excess lipids are redirected at the organismal level toward thermogenesis and hepatic storage in adults. This process is altogether independent from inflammation and insulin resistance promoted by *Ccr2*-dependent monocytes/macrophages. Our data identify a conserved macrophage-dependent mechanism that controls energy storage, conducive to the design of pharmacological interventions.

## Introductory paragraph

Daily and seasonal variation in caloric intake results in cycles of energy storage and expenditure(*1*). To accommodate these cycles, metazoans have evolved specialized fat storing tissues dedicated to dynamic storage of energy(*1–4*) without the common lipotoxicity associated with fat accumulation in other cells(*1, 2, 4*). These adipose tissues consist of lipid-storing adipocytes, stromal cells, and immune cells such as macrophages. Landmark studies have firmly established that the recruitment of monocytes into adipose and peripheral tissues of obese mice promote inflammation, ectopic fat deposition in the liver, and insulin resistance, via the production of inflammatory cytokines such as TNF by the monocyte-derived macrophages(*5–20*). Genetic or pharmacological inhibition of the *Ccr2*-dependent recruitment of bone marrow-derived monocytes into tissues(*21*) prevents or alleviate the metabolic syndrome in obese animals and *Ccr2*-dependent macrophages are a therapeutic target in metabolic diseases associated with obesity(*11, 15–18, 22*). Although CCR2 blockade or deficiency does not prevent weight gain and obesity itself, but only its inflammatory and metabolic complications, recent studies in new models of macrophage-deficient mice suggest that macrophages may regulate mouse weight and adiposity. Trib1-KO mice are lean or even lipo-dystrophic and develop a metabolic syndrome(*23*). In contrast, CSF1R blockade keeps mice lean under a high-fat diet and also protects them against the metabolic syndrome(*24, 25*). Finally, *Trem2*-KO mice(*26*) develop worse obesity and metabolic syndrome. Mechanisms proposed to account for these phenotypes include the pro-adipogenic role of TRIB1-dependent macrophages(*23*), the control of feeding behavior by microglia regulating MBH neurons sensitivity to leptin(*24, 25*), and the global control by *Trem2*(*26*) of a lipid-associated macrophage program. Although the systemic effects of CSF1R blockade and germline deletion of *Ccr2, Trib1*, or *Trem2* combined with the developmental and functional heterogeneity of macrophages(*3, 6, 23, 26–47*) may confound the analysis of macrophage functions in these models, these data nevertheless strongly suggest that macrophages are involved, directly or indirectly, in the regulation of energy storage. In this study we therefore attempted to identify a molecular mechanism that mediate the direct control of adiposity and energy storage by macrophages.

## Results

### Csf1r-dependent lipid storage in newborn, hyperphagic, or high-fat diet fed adult mice

As indicated above 6 week-old adult wild-type C57/Bl6 mice fed a high-fat diet (45% kCal from fat, 4.73 kCal/g) and treated with the CSF1R tyrosine kinase inhibitor PLX5622 (PLX, 1,200 mg PLX5622/kg of food) for 8 weeks stayed lean, with fat pads half the weight of controls (**Figure 1A, Fig. S1**), contrary to *Ccr2*-deficient mice which become obese (*11*)(**Figure 1A, Fig. S2**). In contrast to *Ccr2*-deficient mice, white adipocytes from PLX-treated mice did not increase their Bodipy^+^ lipid content thus lacking the hypertrophy normally induced by high-fat diet (**Figure 1B,C**). Mice treated with a blocking anti-CSF1R antibody (AFS98, 50 mg/kg, i.p., thrice a week (*48*)) presented with a similar phenotype (**Figure 1A, Fig. S1**), indicating that an off-target effect of the kinase inhibitor is unlikely. PLX treatment also reduced obesity and adipocyte hypertrophy in hyperphagic leptin-receptor deficient mice (*Lepr^−/−^, db/db*) fed a control diet (10% kCal from fat, 3.85 kCal/g) (**Figure 1A,B, Fig. S1**). These data altogether indicate that CSF1R blockade prevents fat storage and adipocyte hypertrophy in high fat diet-fed and hyperphagic mice, in a *Ccr2*- and, at least in part, leptin-independent manner.

**Figure 1.**
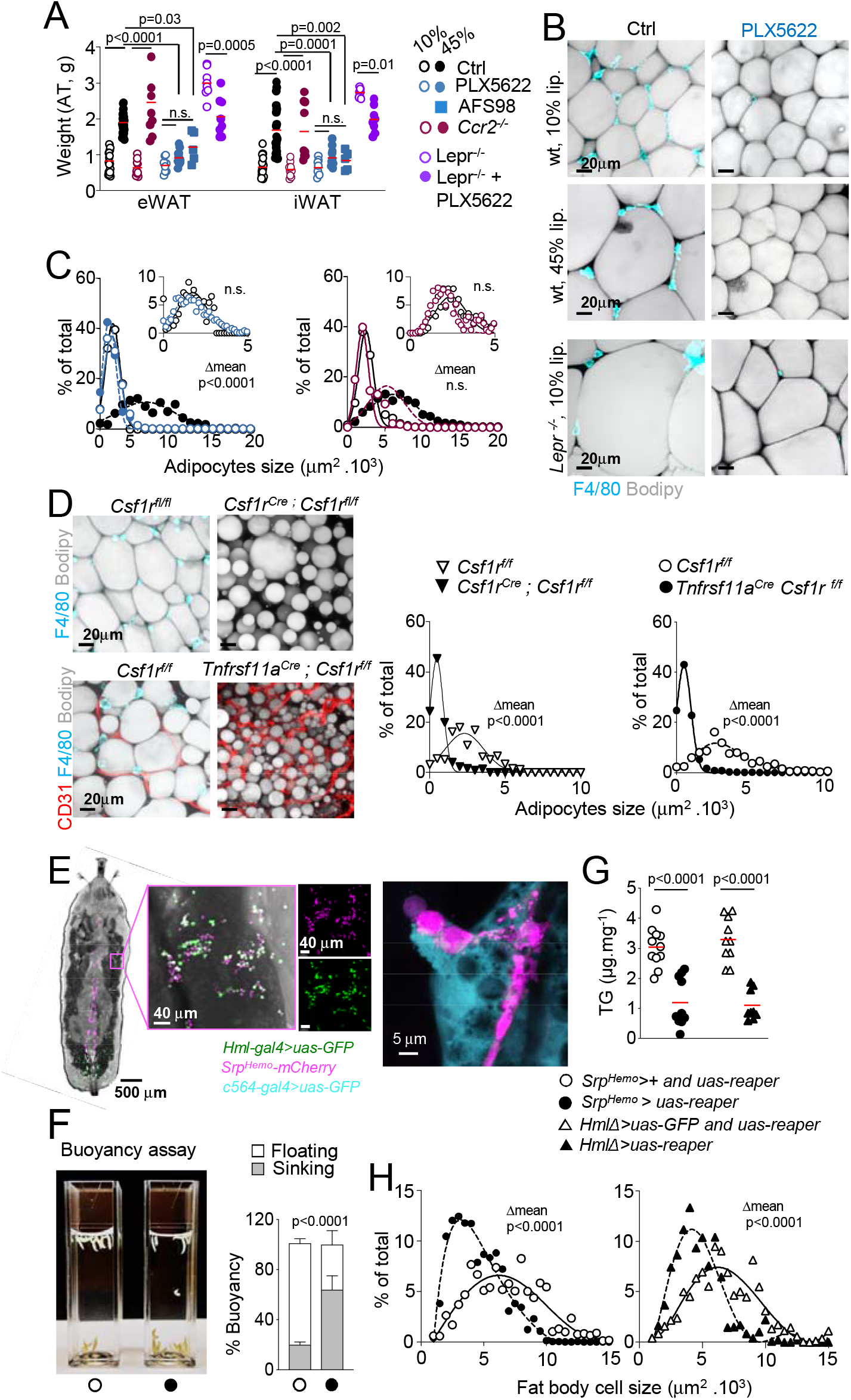
Ccr2-independent macrophages in mice and hemocytes in Drosophila control lipid storage in adipocyte and fat body cells, respectively. (A) Weight of subcutaneous (iWAT) and visceral (eWAT) fat-pads from 14-week-old C57/Bl6 mice, *Ccr2^−/−^* mice, and 12-week-old *Lepr^−/−^* mice all subjected to indicated diets and treatments for 8 weeks. n= 12 to 20 mice per experimental group, from 4 independent experiments for C57/Bl6 mice, and n= 8 to 9 mice per experimental group, from 2 independent experiments for *Ccr2^−/−^* mice, and *Lepr^−/−^* mice. Dots represent individual mice. Statistics: *p* values are obtained by comparing mean weight using one-way ANOVA with Sidak’s correction for multiple group comparison. (B) Representative fluorescent whole mount images stained for F4/80 and Bodipy for inguinal white adipose tissues (iWAT) from 14-week-old C57/Bl6 mice and 12-week-old *Lepr^−/−^* (db/db) mice, all subjected to indicated diets and treatments for 8 weeks. (C) Quantitative analysis of adipocytes size in fat pads from C57/Bl6 and *Ccr2^−/−^* mice from (A,B). n=10 C57/Bl6 and n=6 *Ccr2^−/−^* mice per experimental group, from 2 to 3 independent experiment. 100-200 adipocytes per fat-pad were measured using bitplane Imaris image analysis software in 3 different areas. Dots represent % of adipocytes per size intervals. Statistics: p values are obtained by comparing mean adipocyte sizes using one-way ANOVA with Sidak’s correction for multiple group comparison. See also supplemental figure 1 and 2. (D) Representative fluorescent whole mount images stained for F4/80, Bodipy and CD31 (Left) and quantitative analysis of adipocytes size (Right) for inguinal white adipose tissues (iWAT) from 4-week-old *Csf1r^Cre^; Csf1r^f/f^* mice, *Tnfrsf11a^Cre^; Csf1r^f/f^* mice and their control littermates. Fat pads from 6 mice, obtained from 3 to 6 litters, are analyzed for each group and 100-200 adipocytes were measured using bitplane Imaris image analysis software in 3 different areas per fat pad. Dots represent % of adipocytes per size intervals. Statistics: p values are obtained by comparing mean adipocyte sizes using t-test. See also supplemental figure 3 and 4. (E) Representative fluorescent images of L3 larvae depicting the expression pattern of hemocyte-specific drivers Hemolectin (Hml, *Hml-gal4>uas-GFP*) and Serpent (Srp, *Srp^Hemo^_mCherry*) and the contact between hemocytes and fat-body cells (*c564-gal4>uas-GFP*). (F) Quantitative analysis of buoyancy in wandering L3 larvae from *Srp^Hemo^-gal4>uas-reaper* and control lines from n=6 independent experiments from 3 crosses, each with experimental groups of 20 larvae. Statistics: Buoyancy (mean ± SD) is compared between groups using t-test. (G) Quantification of triglyceride level (TG) normalized to total protein measured in *Srp^Hemo^_gal4>uas-reaper* L3 larvae and controls in n=11 independent experiments from 4 crosses, each with experimental groups of 10 larvae, and in *Hml-gal4>uas-reaper* and controls in n=10 independent experiments, from 4 crosses, each with experimental groups of 10 larvae. Dots represent experimental groups. Statistics: Mean ± SD TG level are compared between genotypes using t-test. (H) Quantitative analysis of Bodipy^+^ fat body cells size in wandering L3 *Srp^Hemo^-gal4>uas-reaper* and controls from n=6 experiments from 3 crosses, each with experimental groups of 20 larvae, and *Hml-gal4>uas-reaper* and controls from n=9 experiments from 4 crosses, each with experimental groups of 20 larvae. For each replicate 100 fat body cells were measured using bitplane Imaris image analysis software. Dots represent % of adipocytes per size intervals. Statistics: p values are obtained by comparing mean fat body cells size between genotypes using t-test.

The initial hypertrophy of adipocytes, which allows for fat storage during the first month of life, was abolished by global genetic deletion of *Csf1r* in *Csf1r^Cre^; Csf1r^f/f^* mice (**Figure 1D, Fig. S3**). Similarly, genetic deletion of *Csf1r* or *Pu.1* in resident macrophages (*49, 50*) (in *Tnfrsf11a^Cre^ Csf1r^f/f^* mice and *Tnfrsf11a^Cre^; Pu.1^f/f^* mice) resulted in mice with small fat pads, corresponding to small Perilipin^+^ adipocytes with little Bodipy^+^ lipid content (**Figure 1D, Fig. S4**). In contrast, adipose tissue weight and morphology of adipocytes were normal in young *Ccr2-*deficient mice (**Fig. S2**), as well as in mice carrying a *Csf1r* deletion in the *Flt3*-expressing hematopoietic stem cell lineage (*50, 51*) (**Fig. S2**). These data suggest that in addition to limiting adipocyte hypertrophy in adult hyperphagic and high-fat diet fed mice, CSF1R deficiency in the *Ccr2-* independent resident macrophage lineage may also prevent the initial hypertrophy of adipocytes in newborn mice, although the adipocyte phenotype of genetic mutant animals may be compounded by the effects of global macrophage deficiency during development.

### Macrophage-dependent lipid storage in Drosophila larvae

Following-up on the hypothesis that the adipocyte phenotype may be attributable to a macrophage defect, and because fat storing cells and macrophages are conserved across the animal kingdom, we generated macrophage-less *Drosophila* larva by inducing apoptosis in hemocytes (*Drosophila* macrophages) using either *Srp^Hemo^-gal4>uas-reaper* or *Hml-gal4 >uas-reaper* lines (*52–54*) (**Figure 1E-H, Fig. S5**). In both case, we observed reduced buoyancy (*55*) in hemocyte-deficient wandering L3 larvae, corresponding to a ~60% reduction in triglyceride content and fat body cell size as compared to control larvae (**Figure 1F-H**). Genetic labeling of *Drosophila* hemocytes, with *Hml-gal4>uas-GFP* (*56*) or *Srp^Hemo^-mCherry* (*54, 57*) reporters also confirmed a close association between hemocytes and fat body cells, labeled with *c564-gal4>uas-GFP* (*53, 58*) in L3 larvae (**Figure 1E**). These data suggested that a genetic screen for macrophage-derived factors may unveil a conserved mechanism by which *Drosophila* and mouse macrophages may control lipid storage and adipocyte hypertrophy.

### Macrophage-derived PDGF-family growth factor Pvf3 controls lipid storage in Drosophila fat body cells

The *Drosophila* larvae is a highly tractable model to screen for the role of conserved families of growth factors and cytokines produced by macrophages (*56, 59, 60*). Among these, hemocytespecific RNAi of the PDGF family ortholog (*59, 61*) *Pvf3 (Srp^Hemo^-gal4>uas-pvf3-IR*) decreased triglyceride content (**Figure 2A**) and fat body cell size by >50% (**Figure 2B,C, Fig. S5**). Hemocyte-specific RNAi of the closely related *Pvf1*, presented with a milder phenotype (**Figures 2A, B**). To confirm the role of *Pvf3* and control for off-target effects of the RNAi constructs, we analyzed *Pvf3* mutant flies (*Pvf3^EY09531^*, **Figure 2D-F**). These experiments showed that L3 *Pvf3^EY09531^* larvae have reduced buoyancy (**Figure 2D**), a >50% reduction in triglyceride content (**Figure 2E**), and small fat body cells in comparison to controls (**Figures 2F**), while the genetic rescue of *Pvf3* expression in *Pvf3^EY09531^* larvae (*Pvf3^EY09531^; He-gal4>uas-pvf3* (*62*)) restored triglyceride content and fat body cell size (**Figure 2E,F**). In addition, since all *Drosophila* PDGF family orthologs share the same receptor *Pvr* (*59, 61*), which is expressed by fat body cells (*61, 63*), we generated fat body-specific *Pvr* RNAi (*c564-gal4>uas-pvr-IR*), which yielded L3 larvae with reduced triglyceride content and small fat body cells (**Figures 2G,H**). These data strongly suggest that *Drosophila* hemocytes control fat body cell size and triglyceride storage via production of the PDGF-family growth factor *Pvf3* by hemocytes, likely acting on *Pvr* expressing fat body cells.

**Figure 2.**
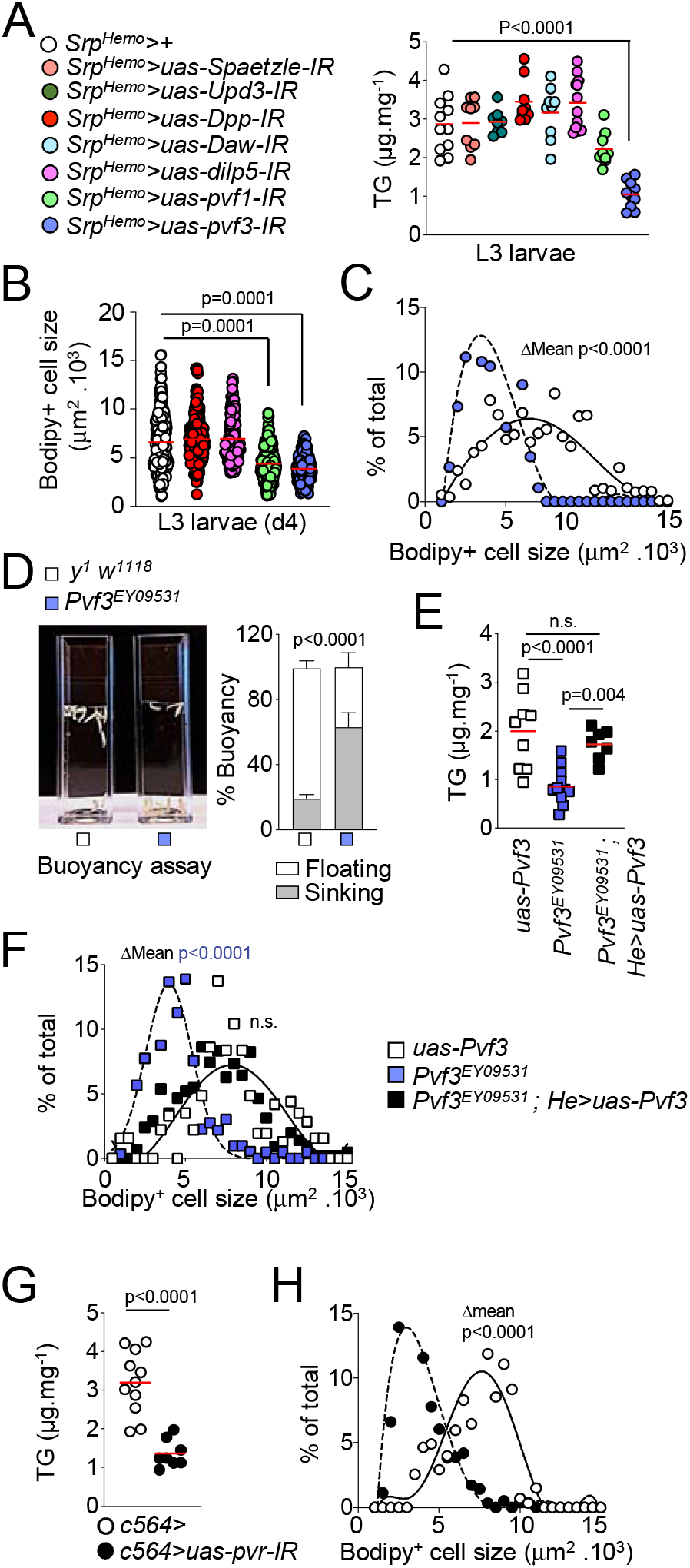
Hemocyte-derived PDGF-family growth factor Pvf3 controls lipid storage in Drosophila fat-body cells. (A) Quantification of triglyceride level (TG) normalized to total protein measured in wandering L3 larvae from flies with hemocyte (macrophage)-specific RNAi for the indicated genes and *Srp^Hemo^*>+ control. For each RNAi, n=8-12 experiments from 4 crosses are analyzed, each with experimental groups of 10 larvae. Dots represent means of individual experiments. *Statistics*: p values are obtained by comparing mean TG levels in mutants and *Srp^Hemo^*>+ control using oneway ANOVA with Dunnett’s correction for multiple group comparison with a single control. (B) Quantification of Bodipy^+^ fat body cell size in L3 larvae from flies with hemocyte-specific RNAi for the indicated genes and *Srp^Hemo^*>+ control. For each RNAi, n=4 to 7 experiments from 3 to 6 crosses were analyzed, each with experimental groups of 10 larvae, and 100 cells per replicate are measured. Dot represents individual fat body cells. *Statistics*: p values are obtained by comparing mean Bodipy^+^ fat body cell size using one-way ANOVA with Dunnett’s correction for multiple group comparison with a single control. (C) Analysis of Bodipy^+^ fat body cells size distribution in L3 *Srp^Hemo^-gal4>uas-pvf3-IR* and control larvae in n=3 experiments from 3 crosses, each with experimental groups of 10 larvae and 100 fat body cells were measured using bitplane Imaris image analysis software for each replicate. Dots represent % of fat-body cells per size intervals. *Statistics*: p values are obtained by comparing mean adipocyte sizes using t-test (D) Quantitative analysis of buoyancy in wandering L3 larvae from *Pvf3^EY09531^* and control *uas-Pvf3* L3 larvae in n=6 independent experiments, from 3 crosses, each with experimental groups of 20 larvae. Statistics: Buoyancy (mean ± SD) is compared between groups using t-test. (E) Quantification of triglyceride level (TG) normalized to total protein measured in *Pvf3^EY09531^*, *uas-Pvf3 (control) and Pvf3^EY09531^; He>uas-Pvf3* (*rescue*) L3 larvae. For each genotype n=7-12 experiments from 4 crosses were analyzed, each with 10 larvae per experimental group. Dots represents individual experimental groups. Statistics: Mean TG levels are compared using oneway ANOVA with Sidak’s correction for multiple group comparison. (F) Analysis of Bodipy^+^ fat body cells size distribution in larvae from (E), in n=5 experiments per genotypes from 3 crosses, each with 10 larvae per experimental group, and 100 fat body cells were examined using bitplane Imaris image analysis software for each replicate. *Statistics*: p values are obtained by comparing mean adipocyte sizes using one-way ANOVA with Sidak’s correction for multiple group comparison as in (E). (G) Quantification of triglyceride level (TG) normalized to total protein measured in L3 larvae from flies with fat body-specific RNAi of Pvr (the Pvf receptor) and controls. For *Pvf* RNAi flies and controls n=8 and n=11 independent experiments respectively were performed, from 4 crosses, and with 10 larvae per experiment. Dots represents individual experimental groups. Statistics: Mean TG levels are compared between the 2 genotypes using t-test. (H) Analysis of Bodipy^+^ fat body cells size distribution in L3 larvae from (G). n=4 experiments from 4 crosses, 10 larvae per experiment, and 100-150 cells per replicate were analyzed using bitplane Imaris image analysis software. Dots represent % of fat-body cells per intervals of size. Statistics: Mean Bodipy^+^ fat body cells size are compared between the 2 genotypes using t-test.

### Macrophage-derived PDGFcc controls lipid storage in newborn and adult mice

Among mouse *Pvf* orthologs, data from the ImmGen Consortium(*64, 65*) indicate that *Pdgfc* is the most abundantly expressed PDGF/VEGF family member in mouse fat-associated macrophages (**Figure 3A**). In addition, we found that *Pdgfc* expression was upregulated in fat tissue of high-fat diet fed wild-type mice and hyperphagic *db/db* mice, in a *Ccr2*-independent but CSF1R-dependent manner (**Figure 3B**). Additionally, *Pdgfc* expression was severely decreased in fat pads from 3-4 week old *Csf1r^Cre^; Csf1r^f/f^* mice (**Figure 3C**). These data support the hypothesis that PDGFcc produced by macrophages might be the growth factor involved in adipocyte hypertrophy in mice. We therefore generated *Csf1r^Cre^; Pdgfc^f/f^* mice (**Figure 3D-G, Fig. S6**). Adipocytes developed normally in *Csf1r^Cre^; Pdgfc^f/f^* mice, based on *Perilipin* and *Ucp1* expression (**Figure 3E,F**), but their lipid content were reduced (**Figure 3E, G**) and white adipose tissue weight was decreased by ~50%, in 4 week-old *Csf1r^Cre^; Pdgfc^f/f^* mice (**Fig. S6**). In contrast to *Csf1r*-deficient mice (**Fig. S3**), *Csf1r^Cre^; Pdgfc^f/f^* mice were not runted or osteopetrotic (**Figure 3D, Fig. S6**), suggesting macrophage functions are otherwise largely intact and therefore the adipocyte phenotype is not compounded by a failure to thrive as is the case in mice with global CSF1R deficiency. To test the hypothesis that PDGFcc also controls lipid storage in adult mice, 6-week-old C57BI/6J mice were treated with anti-PDGFcc neutralizing antibodies (AF1447, 50 μg/mouse, i.p., thrice a week) for 6 weeks (**Figure 3H-J, Fig. S6**). We found that mice treated with anti-PDGFcc under a lipid-rich diet remained lean (**Figure 3H,I**), with small adipocytes (**Figure 3J, Fig. S6**). Altogether, these results very strongly suggest that production of the *Pvf3* ortholog PDGFcc by macrophages controls lipid storage in adipocytes of newborn mice, while its blockade in adult mice prevents adipocyte hypertrophy in response to high fat diet.

**Figure 3.**
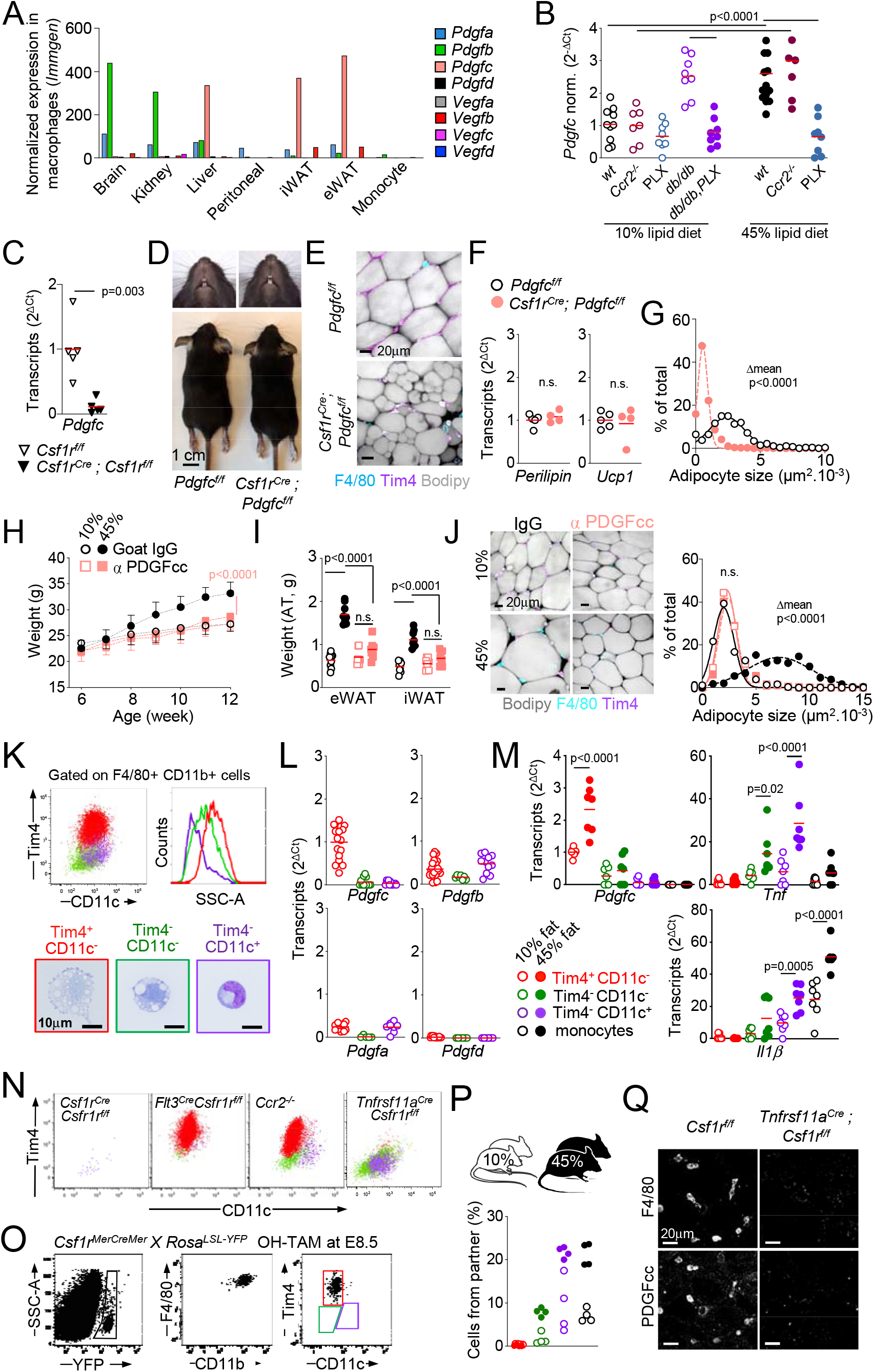
Resident macrophage-derived PDGFcc is required for adipocyte hypertrophy in mice. (A) Normalized expression of *Pdgf/Vegf* family transcripts in murine macrophages from indicated tissues, data are obtained from the Immgen database, and expression values normalized by DESeq2. (B) qPCR analysis of *Pdgfc* transcripts normalized to *Gapdh* (2^−ΔCt^) in inguinal white adipose tissues (iWAT) from 14-week-old C57/Bl6 mice, *Ccr2^−/−^* mice and 12-week-old *Lepr^−/−^* (*db/db*) mice all subjected to indicated diets and treatments for 8 weeks. Fat pads from n= 8 to 15 mice from 2 to 3 independent experiments were analyzed for each group and experimental condition. Dots represent values for individual mice. Statistics: p values were obtained by comparing mean expression values using one-way ANOVA with Sidak’s correction for multiple group comparison. (C) qPCR analysis of *Pdgfc* transcripts, normalized to *Gapdh* (2^−ΔCt^), in inguinal white adipose tissues (iWAT) from 4-week-old *Csf1r^Cre^; Csf1r^f/f^* mice and littermate controls. Fat pads of n= 5 mice per genotype from 3 independent litters were analyzed. Dot represents values for individual mice. Statistics: p values were obtained by comparing mean expression values between different genotypes using t-test. (D,E) Representative photographs (D) and whole mount images of inguinal fat pads stained for F4/80, Tim4, and Bodipy (E) from *Csf1r^Cre^; Pdgfc^f/f^* mice and control littermates at 4 weeks of life. (F) qPCR analysis of *Perilipin* and *Ucp1* transcripts normalized to *Gapdh* (2^−ΔCt^), in inguinal white adipose tissues (iWAT) from 4-week-old *Csf1r^Cre^; Pdgfc^f/f^* mice and littermate controls. Fat pads of n= 4 to 5 mice from 4 litters were analyzed per genotype. Dot represents values for individual mice. Statistics: p values were obtained by comparing mean expression between different genotypes using t-test. (G) Quantitative analysis of adipocytes size for inguinal white adipose tissues (iWAT) from (F). Fat pads from n= 4 mice were analyzed for each genotype. 100 adipocytes were measured per fat pad using bitplane Imaris image analysis software, from 3 different areas. Dots represent % of adipocytes per size interval. Statistics: p values were obtained by comparing mean adipocyte size between different genotypes using t-test. (H) Weight gain of 6-week-old C57Bl/6J mice fed a 10% or 45% lipid diet and treated with anti-PDGFcc antibodies or control goat IgG for 6 weeks. n=9 mice per experimental group, from 2 independent experiments. Mice were weighted weekly for 6 weeks. Dots and bars represent mean body weight ± SD. Statistics: p values were obtained by comparing the mean body weight between experimental groups using two-way ANOVA with Sidak’s correction for multiple group comparison. (I) Weight of subcutaneous (iWAT) and visceral (eWAT) fat-pads from 12-week-old C57/Bl6 mice, from (H). n=8 mice per experimental group, from 2 independent experiments. Dots represent individual mice. Statistics: *p* values are obtained by comparing mean adipose tissue weight using one-way ANOVA with Sidak’s correction for multiple group comparison. (J) Whole mount images and quantitative analysis of adipocytes size for inguinal white adipose tissues (iWAT) from 12-week-old C57/Bl6 mice. Fat pads from n= 9 mice from 2 independent experiments were analyzed for each group and experimental condition and 100 adipocytes were measured using bitplane Imaris image analysis software in 3 different areas per fat pad. Dots represent % of adipocytes per size interval. Statistics: p values were obtained by comparing mean adipocyte sizes using one-way ANOVA with Sidak’s correction for multiple group comparison. (K) Representative tSNE color coded flow plot (Top) and May-Grunwald Giemsa staining (Bottom) of Csf1r-dependent F4/80^+^ macrophages subsets from C57Bl/6J mice iWAT. Fat pads from 30 wild type mice were analyzed with similar results, see supplemental figure 7. (L) qPCR analysis of *Pdgfa, Pdgfb, Pdgfc*, and *Pdgfd* transcripts, normalized to *Gapdh* and *ActinB*, in FACS-sorted iWAT macrophages from 4-week-old *C57Bl/6J* mice. n=10 to 30 sorted macrophage samples from 30 mice were analyzed per group. Each dot represents a sorted sample from an individual mouse. (M) qPCR analysis of *Pdgfc, Tnf*, and *Il1b* transcripts, normalized to *Gapdh* and *ActinB*, in FACS-sorted iWAT macrophages and blood monocytes from 14-week-old C57Bl/6J subjected to the indicated diets and treatments for 8 weeks. n=7 sorted macrophage samples and n=9 blood monocyte samples were analyzed per indicated treatment and diet. Each dot represents a sorted macrophage sample from an individual mouse. Statistics: p values were obtained by comparing mean expression values using one-way ANOVA with Sidak’s correction for multiple group comparison. (N) Representative tSNE color coded flow plot of Csf1r-dependent F4/80^+^ macrophages subsets from the iWAT of *Csf1r^Cre^; Csf1r^f/f^* mice, *Flt3^Cre^; Csf1r^f/f^* mice, *Ccr2^−/−^* mice, and *Tnfrsf11a^Cre^; Csf1r^f/f^* mice. Fat pads from n= 4 to 9 mice per genotype were analyzed from at least 3 independent litters. (O) Analysis of YFP+ embryo-derived macrophages, by flow cytometry of iWAT from 4-week-old *Csf1r^MenCreMer^; Rosa^LSL-YFP^* mice, pulse labeled at E8.5 with 4-hydroxy tamoxifen. Fat pads from 10 mice from 8 independent litters were analyzed with similar results. (P) Analysis of macrophage dynamic in parabiotic pairs, by flow cytometry of blood monocytes and adipose tissue macrophages from iWAT of CD45.1 / CD45.2 female parabiotic pairs maintained for 8 weeks on 10% or 45% fat diet before analysis at 14-week-old. n= 8 mice, each dot represents one mouse. (Q) Fluorescent whole mount images stained for F4/80, PDGFcc, and Bodipy from the iWAT of 4-week-old *Tnfrsf11a^Cre^; Csf1r^f/f^* mice and littermate control mice. Images are representative of n= 5 fat pads from 5 mice per genotype.

### PDGFcc production by resident macrophages in response to high fat diet in adults

Flow cytometry, RNA-seq, and qPCR analyses of myeloid cells within visceral and subcutaneous white adipose tissue indicate the presence of three main *Csf1r*-dependent F4/80^+^ subsets: large Tim4^+^ F4/80^+^ macrophages and two populations of small Tim4^−^ F4/80^+^ cells (**Figure 3K, Fig. S7,8**). *Pdgfc* was preferentially expressed in Tim4^+^ F4/80^+^ macrophages (**Figure 3L**) and was inducible by high-fat diet (**Figure 3M**). In contrast, the production of *Tnf* and *Il1b* was restricted to small Tim4^−^ F4/80^+^ cells and monocytes (**Figure 3M**). Tim4^+^ macrophages were still present in *Ccr2*-deficient and *Flt3^Cre^;Csfr1r^f/f^* (*50, 51*) mice, but absent in *Tnfrsf11a^Cre^;Csfr1^ff^* mice (**Figure 3N, Fig. S2,4**), while Tim4^−^F4/80^+^ monocyte/macrophages are absent or reduced in *Ccr2*-deficient and *Flt3^Cre^;Csfr1r^f/f^* (*50, 51*) mice (**Figure 3N, Fig. S2**). Lineage mapping analysis of CSF1R^+^ erythro-myeloid progenitors (EMP) labelled at embryonic day E8.5 showed post-natal labeling of large adipose tissue F4/80^+^ Tim4^+^ macrophages, but not of small F4/80^+^ Tim4^−^ cells (**Figure 3O, Fig. S7**), and parabiosis experiments showed that Tim4^+^ F4/80^+^ cells do not exchange between parabionts in contrast to F4/80^+^ Tim4^−^ cells and monocytes (**Figure 3P**), altogether indicating that *Pdgfc* is preferentially expressed in embryo-derived resident macrophages. Finally, immunofluorescence analysis confirmed that PDGFcc expression is undetectable in fat-pads of *Tnfrsf11a^Cre^;Csfr1r^f/f^* mice (**Figure 3Q**). Global transcriptional analysis (RNA-seq) further showed that Tim4^+^ and Tim4^−^ cells cluster by population rather than diet in multidimensional scaling and hierarchical clustering of differentially expressed genes (**Fig. S8 and Table S1**). In particular, the analysis of genes associated with M1 or M2 polarization of macrophages (*66*) did not indicate an apparent polarization of macrophage in response to dietary changes (**Fig. S8**). Stable transcriptional signatures associated with Tim4+ resident macrophages primarily included homeostatic processes (Cluster 1, 7, **Fig. S8**), while Tim4^−^ monocyte/macrophages highly expressed genes associated with innate immune response and cell proliferation (Clusters 2 and 5, **Fig. S8**). These data altogether indicate that F4/80^+^ Tim4^+^ resident adipose tissue macrophages (*27–33*) are the main *Pdgf*c-producing cells in mouse fat pads and respond to high-fat diet by increasing *Pdgf*c expression, in contrast to F4/80^+^ Tim4^−^ *Ccr2*-dependent cells which respond to high-fat diet by producing *Tnf* and *111b*.

### PDGFcc blockade results in increased energy expenditure and hepatic storage

To further investigate the consequence of PDGFcc blockade and the mechanisms that underlie leanness at the organismal level, we performed metabolic studies in high-fat diet fed mice. Food intake and fecal caloric density in anti-PDGFcc treated mice were comparable to controls (**Figure 4A,B and Fig. S6**), indicating that PDGFcc blockade does not affect food intake or intestinal absorption, as observed similarly for CSFR1 inhibitor (PLX5622) treated mice (**Fig. S1**). In addition, PDGFcc blockade did not protect mice against high-fat diet induced hepato-steatosis (**Figure 4C**), inflammation, or insulin resistance (**Fig. S6**). This was in sharp contrast to *Ccr2*-deficient mice and PLX-treated mice, which lack *Ccr2*-dependent inflammatory monocytes/macrophages, and are protected against hepato-steatosis, inflammation, and insulin resistance (*11, 15–18, 22*) (**Fig. S1,2**). These data indicate that PDGFcc blockade decreases adiposity, without limiting food intake, intestinal absorption, or the metabolic syndrome, the latter being promoted by a different cell type, i.e. *Ccr2*-dependent monocyte/macrophages.

**Figure 4.**
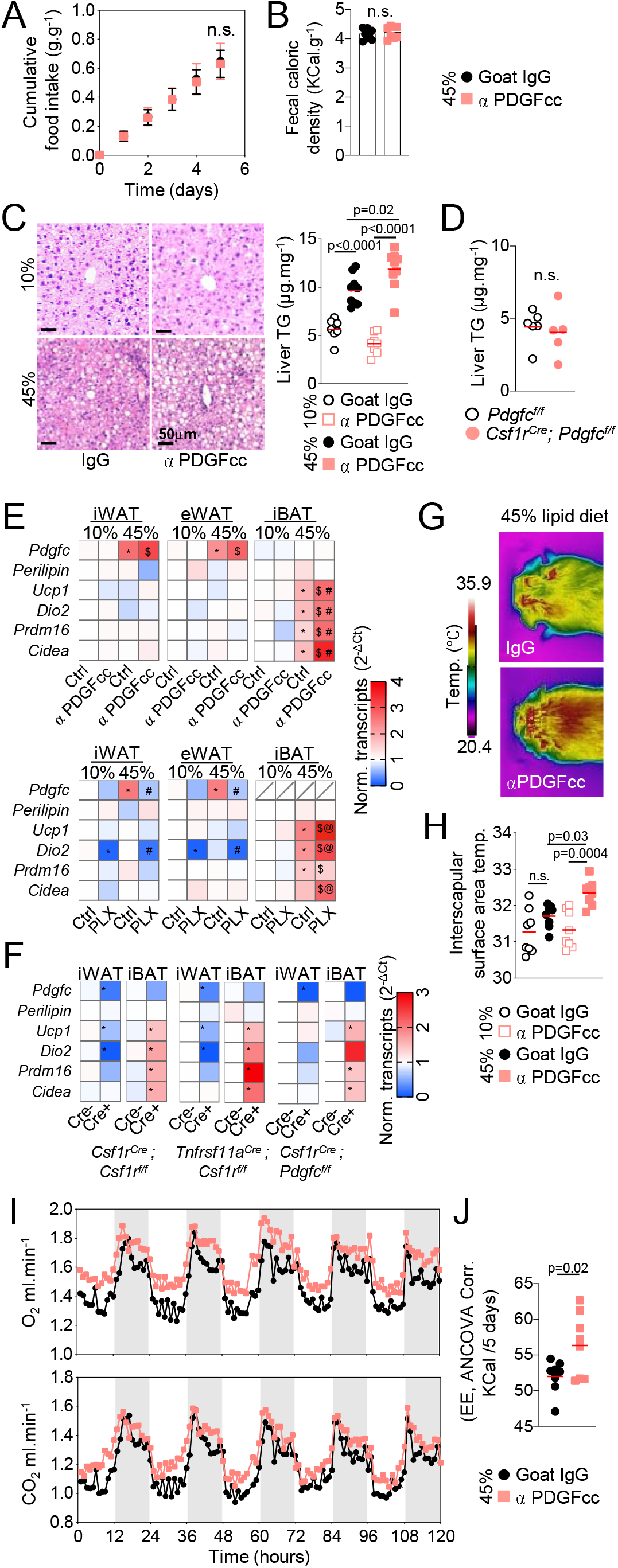
PDGFcc blockade results in increased energy expenditure and hepatic storage. (A, B) Cumulative food intake measured gravimetrically over 5 days (A) and fecal caloric density measured by bomb calorimetry on the last day of the experiment (B) in groups of 8 week-old C57Bl/6J mice fed a 45% lipid diet and treated with anti-PDGFcc antibodies (n=8) or goat IgG (n=8). Dots represent mean cumulative food intake ± SD in (A) and individual mice in (B). Statistics: Mean cumulative food intake and fecal caloric density are compared by t-test. (C) Representative H&E staining (Left) and quantification of triglycerides (TG) content per mg of tissue (right) of livers from 12-week-old C57Bl/6J mice (from Figure 3H-J) fed a 10% or 45% lipid diet for 6 weeks and injected with anti-PDGFcc antibodies or with Goat IgG. n= 8 mice per group, from 2 independent experiments. Dots represent individual mice. Statistics: mean and SD of experimental groups are compared by one-way ANOVA with Sidak’s correction for multiple group comparison. (D) Quantification of triglycerides (TG) content per mg of tissue from 4-week old *Csf1r^Cre^; Pdgfc^flox/flox^* mice and control littermates. n= 4 and 6 mice respectively from 3 litters. Dots represent individual mice. Statistics: mean and SD of experimental groups are compared by t-test. (E) Heatmap representation of *Pdgfc, Perilipin, Ucp1, Dio2, Prdm16*, and *Cidea* transcripts expression (2^−ΔCt^, relative to *Gapdh*) in iWAT, eWAT, and iBAT from 12-week-old C57Bl/6J mice treated for 6 weeks with anti-PDGFcc antibodies or goat IgG. Color scale is normalized on the mean expression of each transcript in the fat tissues from mice on control diet (10% lipid) treated with control goat IgG. n=5-10 mice per group. Statistics: mean expressions values are compared by one-way ANOVA with Sidak’s correction for multiple group comparison * indicates p<0.01 as compared to 10%/goatIgG controls, $ indicates p<0.01 as compared to 10% /? PDGFcc treated mice, # indicates p<0.001 as compared to 45%/ goat IgG controls. (F) Heatmap representation as in (E), from iWAT, eWAT, and iBAT from 14-week-old C57Bl/6J mice treated for 8 weeks with PLX5622 or vehicle control. n=5-10 mice per group. Color scale is normalized as in (E) on 10% lipid diet/ vehicle control. Statistics: As in (E), * indicates p<0.01 as compared to 10%/vehicle controls, $ indicates p<0.01 as compared to 10% PLX-treated mice, @ indicates p<0.01 as compared to 45%/vehicle controls. (G) Heatmap representation as in (E) from iWAT and iBAT from 4-week-old *Csf1r^cre^; Csf1r^f/f^* n=5 mice, *Tnfrsf11a^cre^; Csf1r^f/f^*, n=12 mice*, Csf1r^Cre^; Pdgfc^f/f^*, n=5 mice, and their littermate controls, n=5-12 mice per group. Color scale is normalized on Cre-negative controls. Statistics: mean expression values are compared by t-test. * indicates p<0.01 as compared to littermate controls. (H, I) Representative infrared images (H) and quantitative analysis of temperature of the top 10% surface area warmest pixels (I) of the interscapular area (iBAT) on day 6 of treatment of C57Bl/6J mice fed a 45% lipid diet and injected with anti-PDGFcc antibodies (n=8 mice) or goat IgG (n=8). Dots represent individual mice. Statistics: mean of experimental groups are compared by one-way ANOVA with Sidak’s correction for multiple group comparison. (J, K) Analysis of Oxygen uptake (VO2) and (VCO2) kinetics (J) and covariance corrected cumulative energy expenditure (EE, ANVOVA Corr.) (K) of C57Bl/6J mice fed a 45% fat diet and injected with anti-PDGFcc antibodies (n=8) or goat IgG (n=8) over a 5 days period in a Promethion Metabolic Screening System. n= 8 mice per experimental group. Dots in J represent mean values over time, and dots in K represent individual mice. Statistics: mean of experimental groups in K are compared using t-test.

6 weeks of treatment with anti-PDGFcc under high fat diet resulted in a 20% increase in ectopic storage of triglycerides in the liver as compared to controls (**Figure 4C**), although this was not observed in developing *Csf1r^Cre^; Pdgfc^f/f^* mice (**Figure 4D**), suggesting that ectopic storage may not be the cause for reduced adipocyte storage. Indeed, although thermogenesis associated transcripts(*67, 68*) were not altered in the white adipose tissue (**Figure 4E,F**), they were induced in the brown fat by high-fat diet, and further increased by PDGFcc blockade and CSFR1 inhibition (**Figure 4E**). Thermogenic transcripts were also induced in the brown fat from *Csf1r^Cre^; Pdgfc^f/f^* mice as well as *Csf1r^Cre^; Csf1r^f/f^* and *Tnfrsf11a^Cre^ Csf1r^f/f^ mice* (**Figure 4F**). Accordingly, PDGFcc blockade raised surface body temperature in the inter-scapular area (**Figure 4G,H**) and increased total energy expenditure (**Figure 4I,J**). These data altogether indicate that reduced lipid storage in white adipocytes is associated with increased energy expenditure at the organismal level, at least in part in the form of thermogenesis by the brown adipose tissue, and with redistribution of excess lipids to the liver in conditions of high-fat diet. Thus, we reasoned that PDGFcc may act either locally by promoting storage by adipocytes in a paracrine manner, and/or systemically by limiting expenditure/thermogenesis and lipid redistribution at the organismal level.

### PDGFcc is required and sufficient for adipocyte hypertrophy in fat-pad explants

We therefore performed loss-of-function and rescue experiments in murine isolated fat-pad explants. Wild-type neonatal (P4) murine epidydimal fat pad *anlagen* contain PDGFRα^+^ cells, capillaries and macrophages, but lack perilipin^+^, bodipy^+^ or LipidTox^+^ adipocytes (**Fig. S9**). Perilipin^+^ bodipy^+^ adipocytes develop in these explants cultured *ex vivo* for 10 days in conventional non-adipogenic medium (*36*) (**Figure 5A, Fig. S9**). We found that treatment of wild-type *anlagen* explants with anti-PDGFcc neutralizing antibodies (AF1447) resulted in the development of smaller Perilipin^+^ *Pparg^+^* cells with scant Bodipy^+^ content, in comparison with *anlagen* treated with control goat IgG (**Figures 5A-D**). In addition, *anlagen* from *Tnfrsf11a^Cre^; Csf1r^f/f^* mice also gave rise to small Perilipin^+^ *Pparg^+^* cells that contain little to no lipids as evidenced by scant Bodipy staining as compared control littermates (**Figure 5E-I**). Moreover, addition of recombinant PDGFcc to *Tnfrsf11a^Cre^; Csf1r^f/f^* fat-pads *anlagen* rescued Bodipy staining and size of adipocytes to near wild-type levels (**Figure 5G-I**). These data demonstrate that PDGFcc produced by macrophages is required and sufficient for lipid storage *ex-vivo* in isolated fat pads explants. PDGFcc appears to promote lipid storage at least in part via a paracrine effect on white adipocytes within fat pads (**Figure 5J**). Therefore, increased energy expenditure *in vivo* is likely to be secondary to the increased systemic availability of lipids that are not stored, although our data do not exclude that PDGFcc may also act systemically, for example on brown adipocytes to alter thermogenesis.

**Figure 5.**
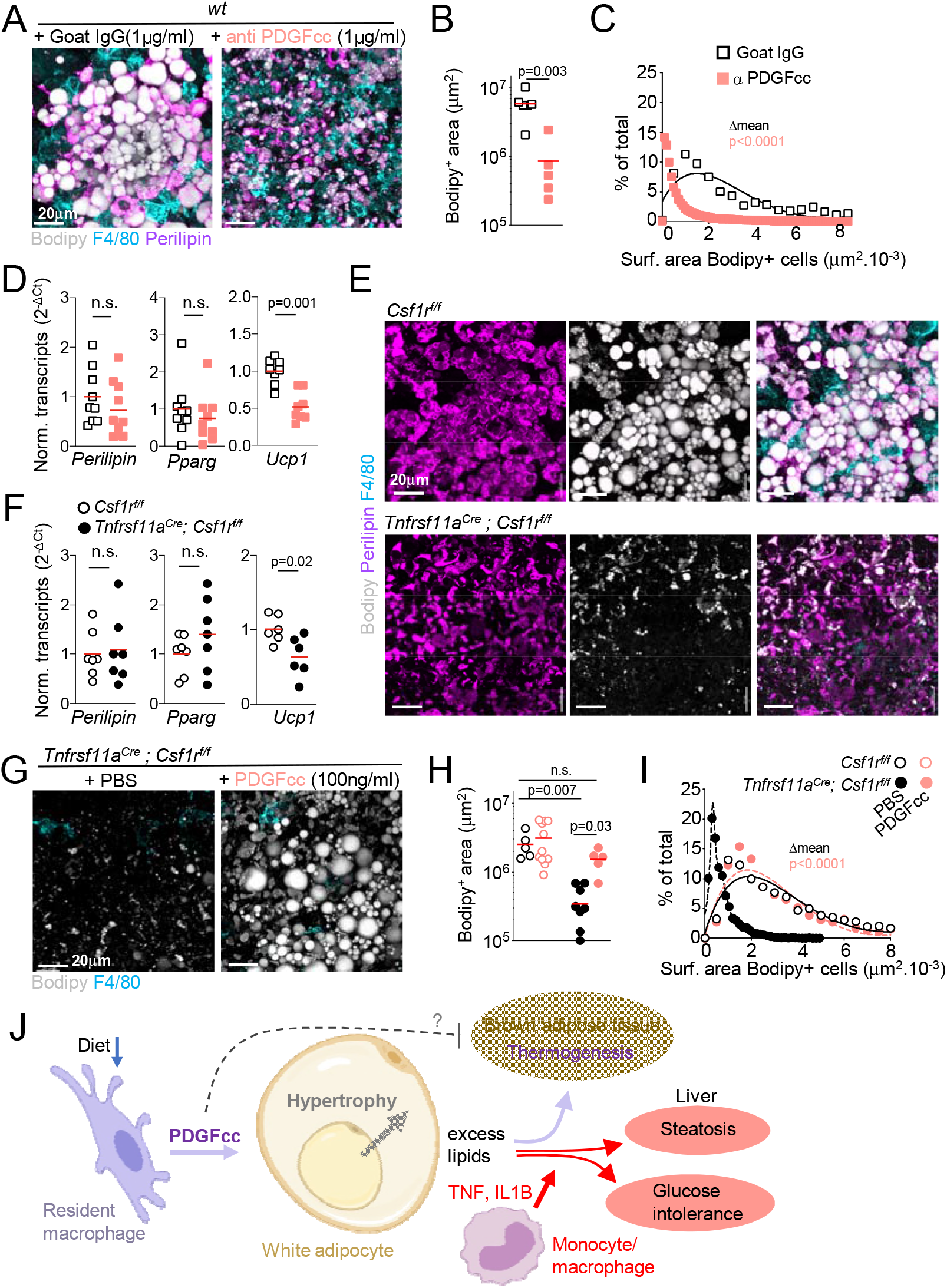
PDGFcc is both required and sufficient for adipocyte hypertrophy in fat tissue explants. (A-C) Representative whole mount staining (A), quantitative analysis of Bodipy^+^ area (B), and quantitative analysis of adipocyte size (C), in wild type eWAT explants cultivated for 10 days (see Extended data and Methods) with Goat IgG (n=5) or anti-PDGFcc neutralizing antibodies (n=5) and stained for Perilipin, F4/80, and Bodipy. Dots represent individual explants in (B), and the % of adipocytes per size intervals in (C). For adipocyte size distribution, >10,000 cells were analyzed per explant using bitplane Imaris image analysis software. Statistics: Mean Bodipy^+^ area in (B) and adipocyte size in (C) were compared between controls and anti-PDGFcc treated explants using t-test. (D) qPCR analysis of *Perilipin, Pparg* and *Ucp1* expression (2^−ΔCt^ calculated relative to *Gapdh*) in wild type eWAT explants cultivated as in A. n=8 to 9 explants per experimental condition, from 2 independent experiments. Dots represent individual explants. Statistics: Results from controls and anti-PDGFcc treated explants were compared using t-test. (E, F) Representative whole mount staining (E) and qPCR analysis of *Perilipin, Pparg* and *Ucp1* expression (2^−ΔCt^ calculated relative to *Gapdh*) (F) of eWAT explants from *Tnfrsf11a^Cre^; Csf1r^f/f^* and *Csf1^f/f^* littermate control mice, from n= 10 and 15 mice respectively, from 5 litters, cultured for 10 days *ex vivo* and stained for Perilipin, F4/80, and Bodipy. Dots represent individual explants. Statistics: qPCR results were compared between mutant and controls littermates using t-test. (G) Representative whole mount staining of eWAT explants from *Tnfrsf11a^Cre^; Csf1r^f/f^ mice* cultivated for 10 days (see Extended data and Methods) with vehicle control (PBS, n=8) or 100ng/ml recombinant PDGFcc (n=5) and stained for F4/80 and Bodipy. (H, I) quantitative analysis of Bodipy^+^ area (H), and quantitative analysis of adipocyte size (I) using bitplane Imaris image analysis software, in explants from *Tnfrsf11a^Cre^; Csf1r^f/f^* mice and *Csf1r^f/f^* littermate controls, cultivated as in (G) with PBS (n= 5 mutants and 8 littermates) or recombinant PDGFcc (n=5 mutants and 11 littermates). Dots represent individual explants in (H), and the % of adipocytes per intervals of size in (I). Statistics: results from genotypes and treatment groups were compared by one-way ANOVA with Sidak’s correction for multiple group comparison. (J) Schematic depicting the distinct roles of resident and bone-marrow-derived macrophages in control of lipid storage and metabolic syndrome.

## Discussion

We report here observations which indicate that PDGF family growth factors (*Pdgfc* and *Pvf3*) produced by resident macrophages in response to dietary changes mediate the storage of lipids in adipocytes of hyperphagic and lipid-rich-diet fed adult mice, developing mice, and *Drosophila* larva. Based on our data, the lean phenotypes observed in the absence of macrophages or PDGFcc is unlikely to be due to decreased food intake, increased leptin signaling, intestinal malabsorption, or white adipose tissue browning. Rather, our data indicate that the diet-regulated production of PDGFcc by macrophages is dispensable for adipocyte differentiation but controls their lipid content in a paracrine manner. Excess lipids which are not stored in the absence of PDGFcc appear to be redirected for thermogenesis or ectopic storage. This function of adipose-tissue resident macrophages, *i.e*. promoting lipid storage via the hypertrophy of specialized fat cells in response to nutrient intake, appears to be evolutionarily conserved in mice and *Drosophila. Drosophila* do not have fibroblasts or endothelial cells, and *Pvr* silencing in fat-body cells phenocopies *Pvf3* silencing in hemocytes, suggesting that the interaction between macrophages and fat cells may be a direct one at least in flies. Nevertheless, we acknowledge that PDGF-receptor signaling in mammals is complex and support many cellular functions(*69–73*), and whether macrophages directly signal to adipocytes in mice, or in this case which member(s) of the PDGF/VEGF- receptor family mediate the effect of PDGFcc on mouse adipocytes are both unknown. Overall, our data suggest that resident macrophages and specialized fat-storing cells constitute a functional unit in metazoans, where macrophages sense the organism’s nutritional state (*74*) and, when intake is high, signal to adipocytes for increased lipid storage through regulated paracrine production of PDGF family growth factors (**Figure 5J**).

Macrophage developmental heterogeneity frequently underlie specialized functions associated with distinct macrophage cell types. In the present case, our data clearly shows that *Ccr2-* independent Tim4^+^ resident macrophages are not sufficient to promote diet-associated inflammation or insulin resistance, while Tim4^−^ *Ccr2*-dependent monocyte/macrophages promote inflammation and the metabolic syndrome (*11, 15–20, 22*), but are neither required nor sufficient for fat storage in white adipose tissue. Therefore our data do not support a model where obesity, inflammation, and insulin resistance in mice would all be controlled by the reprogramming of resident macrophages from a homeostatic into a pro-inflammatory role in the setting of metabolic stress. Instead our data strongly suggest that distinct developmental subsets, *i.e*. resident macrophages and recruited HSC derived monocyte/macrophages, perform distinct functions in the adipose tissue ‘niche’ and are targeted independently by CCR2 and PDGFcc blockade. Altogether, the present study identifies a mechanism that controls adiposity, i.e. the expansion of fat stores, in metazoans in the context of a cellular and molecular ‘dissection’ of macrophage functions, which may help explain and predict the effect of genetic variants and pharmacological interventions on lipid storage.

## Acknowledgements

This study is dedicated to the memory of Lucile Crozet. This work was supported by NIH/NCI P30CA008748 to MSKCC, NIH/NIAID 1R01AI130345, NIH/NHLBI R01HL138090, Ludwig institute for Cancer research basic immunology grant and Cycle for Survival grants to FG. This work was also supported by Leducq transatlantic network of excellence to CKG and FG, NIH/NCI F32CA225036 to NC and Alan and Sandra Gerry Metastasis and Tumor Ecosystems Center fellowship to PLL. The authors thank John Frampton, Jeffrey Pollard, and Thomas Boehm, for providing mouse strains, Harmony Ketchum for her technical assistance with the *Csf1^−/−^* mice, the molecular cytology core at MSKCC for help with preparation of histological samples. The authors acknowledge Maria Pokrovskii and all members of the Geissmann lab for helpful suggestions and editing the manuscript.

## Author contribution

FG, LC, NC designed the study, analyzed data, and wrote the manuscript. LC, NC, and PLL performed and analyzed experiment in mice. NC performed *Drosophila* experiments. EM contributed to generation of *Tnfrsf11a^Cre^; Csf1r^f/f^* mice. TL helped with analysis of flow cytometry data. IRH, CKG analyzed RNAseq experiments and helped preparing the manuscript. PLL helped with immunofluorescence and flow cytometry analyses. RS provided intellectual input. EM present address: Developmental Biology of the Innate Immune System, LIMES Institute, University of Bonn, 53115 Bonn, Germany.

## Notes

### Competing Interest Statement

The authors have declared no competing interest.

