## Supplementary material for "Diet-regulated production of PDGFcc by macrophages controls energy storage": Methods

**Material and methods:**

**Mice and diets.** All experiments on mice were realized in accordance with our animal license issued by the Institutional Review Board (IACUC 15-04-006) from MSKCC. All mice were maintained under SPF conditions. C57Bl/6J male mice were purchased from Jackson laboratories. *Rosa26^LSL-Tomato^* (Stock No: 007908), *Rosa26^LSL-YFP^* (Stock No: 006148), *Rosa26^mTmG^* (Stock No: 007576) and *Lepr ^-/-^(db/db)* (Herein referred to as *Lepr^-/-^*; Stock No: 000697) mice were purchased from Jackson laboratories and bred in house. *Csf1r^Cre^*, *Csf1r^MeriCreMer^*, and *Csf1r^f/f^* mice were obtained from J W. Pollard (The university of Edinburgh), *Cx3cr1^GFP/+^* mice were obtained from Dan Littman (NYU Skirball institute), *Flt3^Cre^* mice were obtained from Thomas Boehm (Max Planck institute), and *Tnfrsf11a^Cre^* mice obtained from Y. Kobayashi (Matsumoto Dental University), all were previously described(*1-4*)*. Ccr2^-/-^* mice were obtained from I F. Charro (UCSF). *Pdgfc^f/f^* mice were purchased from *International Knockout Mouse Consortium*. *Csf1r ^-/-^* mice(*5*) were backcrossed to FVB/NJ for more than 10 generations.

Mice were fed ad libitum a rodent irradiated diet containing 45% kcal from lipids (Research Diets Inc., reference D12451i) or 10% kcal from lipids (Research Diets Inc., reference D12450hi) for 8 weeks. In mice supplemented with the CSF-1R inhibitor PLX5622 (Plexxikon), the aforementioned diets were impregnated with 1200 mg of inhibitor pr kg of food. Mice were weighed thrice per week and then the values were average for the week. *Tnfrsf11a^Cre^; Csf1r^f/f^ , Tnfrsf11a^Cre^; PU.1^f/f^ ,* and *Csf1r^Cre^; Csf1r^f/f^* mice are osteopetrotic and as such lack teeth. To prevent confounding systemic effects due to nutritional intake problems, the *Tnfrsf11a^Cre^; Tnfrsf11a^Cre^; PU.1^f/f^ , Csf1r^f/f^*, *Csf1r^Cre^; Csf1r^f/f^*, and control littermates were fed a nutritionally fortified water gel. The nutritional gels were replaced every 12 hours.

For tissue collection, mice were sacrificed by overdose of anesthetic, with intraperitoneal injection of ketamine (50 mg/kg), xylazine (10 mg/kg) and acepromazine (1.7 mg/kg). Once the withdrawal reflexes (paw) were abolished, intracardiac puncture was performed for blood collection, with an EDTA (100mM, Sigma) coated syringe (26G needle, 1mL syringe, BD). The heart was then perfused with 10mL PBS at room temperature. The mice were then sacrificed by cervical dislocation and the tissues collected.

**Parabiosis and Fate mapping**. For parabiosis experiments, 6 weeks old congenic CD45.1 and CD45.2 were surgically joined(*6*). Skin was incised from elbow to knee, forelimbs and hind limbs were joined with nylon suture, then skin incisions were sutured with stainless steel clips. The mice were fed a sulfatrim diet for 2 weeks following the surgery and before starting 10% lipid diet or 45% lipid diet for 8 weeks. Chimerism in Tim4+ and Tim4- macrophage populations, as well as in liver Kupffer cells and blood lymphocytes and monocytes was assessed by flow cytometry.

Fate mapping experiments in *Csf1r^MeriCreMer^* mice were performed as described (*4*); in brief *Csf1r^MeriCreMer^* females were crossed to *Rosa26^LSL-tomato^ or Rosa26^LSL-YFP^* males and injected intraperitoneally at 8.5 days post coitum with 75mg per kg of body weight of 4-hydroxytamoxifen (Sigma) plus 37.5mg progesterone (Sigma) per kg of body weight to counteract possible abortion induction. Tomato or YFP expression was then assessed in adult progeny by flow cytometry.

**Glucose and insulin tolerance test.** Glucose and insulin tolerance test were performed as described(*7*); in brief, for the glucose tolerance test, the mice were fasted for 4h before experiment, but had water ad libitum. The weight and glycaemia were measured, and the mice were injected intra-peritoneally with 2g glucose (Thermo Scientific) per kg of body weight. Following the glucose injection, glycemia was measured every 30 min for 2 hours. For the insulin tolerance test, the mice were fasted for 4h before experiment, but had water ad libitum. The weight and the glycaemia were measured, and the mice were injected with 0.25U insulin (Sigma) per kg of body weight. Following the glucose injection, glycemia was measured every 30 min for 2 hours. To measure the glycaemia, the tail of the animals was pricked with a 27G needle and a drop of blood was placed on the glucometer (ACCU-CHECK AVIVA, Roche).

**Mice antibody treatment.** Anti-CSF1R (clone AFS98, BioXcell) was administered intraperitoneally at 50 mg/kg in 100µL PBS, and anti-PDGFcc (AF1447 R&D system) at 50 µg/mouse in 100µL PBS. Antibodies were administrated every 2 days from the start of high fat diet. Mice treated with anti-CSF1R antibody also received an injection 2 days prior to high fat diet.

**Metabolic analysis of Leptin receptor deficient mice, PLX5622-treated mice, anti-PDGFcc treated mice, and control C57Bl/6J mice.** Animals were individually housed in a temperature controlled Promethion Metabolic Screening System (Sable Systems International, NV). Food intake was acquired gravimetrically and ambulatory activity was acquired using a laser matrix. Mice were acclimated to this environment on a 12 hr light/dark cycle for 48 hr before the indicated length of recording (24hr to 120hr) period began. Fecal bomb calorimetry was performed on feces collected during the last day of metabolic cage recording period, dehydrated in an oven at 60C for 48 hr, then combusted in technical duplicates with a Parr 6725 Semimicro Calorimeter to determine gross caloric intake.

**Infrared imaging of mice.** Infrared images were taken on the last day of metabolic analysis using a FLIR T430sc Infrared Camera. These images were into RAW files and then analyzed in AMIDE(*8*). A box was drawn over region of interest (ROI); interscapular (70x30), inguinal (35x35) and dorsal. The 10% top warmest pixels were used for interscapular and inguinal quantifications. The 20% top warmest pixels were used for dorsal quantifications. The variance refers to the variance of the voxels in the ROI.

**Triglyceride measurement for mice.** To measure triglycerides, ~ 100 mg of mouse liver was homogenized in 100 μl of PBS + 0.05% Tween 20. Triglyceride was then measured using the free glycerol reagent (Sigma; F6428) as described in the manufacturer’s protocol and normalized to the liver weight.

**Cell suspension preparation for flow cytometr­­­­y and flow cytometry analysis.** For the blood, red blood cells were lysed twice in ACK lysis buffer. Adipose tissue was collected in PBS, incubated for 20 min at 37°C in collagenase II at 2mg/mL (Sigma) in PBS supplemented with 0.25% BSA (Thermo Scientific) and 5mM Cacl_2_ (Sigma) under agitation, before mechanical disruption with a 10mL pipette. Cell suspensions were centrifuged for 10min at 500g(*9*). Liver, brain, and kidney samples were digested for 30 min at 37°C in PBS containing 1mg/ml of collagenase D (Roche), 100 U/ml DNaseI (Sigma), 2.4mg/ml of dispase (Invitrogen) and 3% fetal calf serum (FCS, Invitrogen). Once processed, all samples were resuspended in FACS buffer (PBS, 0.5% BSA and 2 mm EDTA) containing anti-mouse CD16/32 (FcRIII/II, Biolegend Cat#:101302) and anti-mouse CD16.2 (Biolegend Cat#:149502) for 10 min and stained with antibody mixes for 30min on ice. The list of antibodies used can be found in **Table 2**. After 2 washes with FACS buffer, samples were incubated with 2 μM Hoechst 33342 (Thermo Scientific) just prior to analysis using LSR Fortessa X-20 (BD bioscience) or sorting on an ARIA III (BD bioscience). The number of cells per gram of tissue was determined using a cell counter (GUAVA easyCyte HT).

Flow cytometry data were analyzed with Flow Jo 9.9. For t-SNE analysis of the adipose tissue stromal vascular fraction, FCS files of different conditions or genotypes were concatenated. t-SNE algorithm, from Flow Jo 9.9, was used on singlets (determined by FSC-A, SSC-A gate, live cells and FSC-W, FSC-A) with 1000 iterations, perplexity at 20, and Theta at 0.5. All channels with antibody staining were considered as well as FSC-A and SSC-A. Expression of the markers expressed by each cluster was performed after the separation of the concatenated samples.

**Preparation of libraries for RNA sequencing and analysis.** To optimize cell viability, decrease cell activation and improve library quality, a low input RNA sequencing strategy was adopted. Libraries were prepared on 200 cells, reducing the sorting time, and flavopiridol (Sigma) was added during the sample preparation to inhibit transcription.

Blood was processed as described above, with addition, in the antibody mix, of 2µmol/L flavopiridol. For adipose tissue the enzymatic digestion was performed with 4mg/mL collagenase II, resuspended in 0.25% BSA and 5mM CaCl_2_ and supplemented with 2µ mol/L flavopiridol, for 30 min at room temperature under agitation. The rest of the procedure was performed as described above. 200 cells from each sample were directly sorted in a 96 well plate (Biorad) in 4 μL H_2_O containing 0.2% Triton X-100 (Sigma) and 0.8U/mL RNase inhibitor (Clontech). RNA was extracted with RNeasy mini kit (Qiagen), following manufacturer instructions and was quantified with ribogreen quantification (Thermo Scientific). Agilent bioAnalyzer was used for quality control. For each sample, 400pg of were amplified for 14 cycles with SMART-seq V4 (Clonetech) ultra-low input RNA kit for sequencing. Illumina HiSeq libraries were prepared with 10 ng amplified cDNA, using 8 cycles of PCR with Kapa library preparation chemistry kit (Kapa Biosystems). Barcoded samples were run on a HiSeq 2500 1T in a 50bp/50bp paired end run using TrueSeq SBS Kit V3 (Illumina). An average of 33 millions of read was generated per sample, with an average of 57% mRNA bases.

Fastq read files were aligned with Star Version 020201(*10*) to mm10 reference genome and quantified with Homer version 4.9(*11*), then subsequently analyzed in R with Bioconductor package Limma version 3.28.21 and Voom(*12-15*). For the cell-type analysis between groups differential gene expression was performed with FDR corrected P-value < 0.05 and absolute logFC > 1 of any group and the mean expression. K-means clustering was applied with 20 clusters, and highly correlating clusters with Spearman rank correlation higher than 0.9 were subsequently combined resulting in 8 clusters. Clusters were reordered according to number of genes. Functional enrichment was performed using Metascape(*16*). For the effects of food intake, linear model contrasts were set-up for each cell type for pairwise comparisons of the 10% lipid diet, 45% lipid diet, and 45%>10% lipid diet. Explained variance for all expressed genes in the RNA-seq data was calculated using the VariancePartition Bioconductor package(*17*).

**qRT-PCR on sorted cells and adipose tissue samples.** For qRT-PCR, 10000 cells were sorted in 300 μL RNA lysis buffer (Macherey-Nagel, Nucleospin TriPrep), after preparation of the samples as described for RNA sequencing. RNA extraction was performed following manufacturer’s instructions (Macherey-Nagel, Nucleospin TriPrep), RNA concentration was measured with nanodrop2000. cDNA preparation was performed with Quantitect Reverse transcription kit (Qiagen) as per manufacturer’s instructions. qRT-PCR were done with 1 ng cDNA.

For qRT-PCR performed on eWAT and iWAT from *Ccr2^-/-^* and littermate controls as well as *Tnfrsf11a^Cre^; Csf1r^fl/fl^* and littermates, tissue samples were weighed and snap frozen in liquid nitrogen. RNA extraction was performed on 20mg of tissue following the same protocol as for the sorted cells and the qRT-PCR was performed on 10ng cDNA. qRT-PCR are performed on a Quant Studio 6 Flex using TaqMan Fast Advance Mastermix, and TaqMan probes for *ActinB* (Mm02619580_g1), *Gapdh* (Mm99999915_g1), *Tnf* (Mm00443258_m1), Il1b (Mm00434228_m1), *Pdgfa* (Mm01205760_m1), *Pdgfb* (Mm00440677_m1), *Pdgfc* (Mm00480205_m1), *Pdgfd* (Mm00546829_m1), *Bmp2* (Mm01340178_m1), *Pparg* (Mm00440940_m1), *Perilipin* (Mm00558672_m1), *Ucp1* (Mm01244861_m1), *Dio2* (Mm00515664_m1), *Prdm16* (Mm00712556_m1), and *Cidea* (Mm00432554_m1).

**Whole mount imaging and cytology of sorted macrophages.** For whole mount immunofluorescence imaging, after anesthesia and blood collection by cardiac puncture, the mice were perfused with 10 mL PBS at room temperature. Approximately 3 mm^3^ pieces of the tissue were incubated in 4% paraformaldehyde (PFA) (Electron Microscopy) diluted in PBS for 30 min at room temperature with agitation, then rinsed with PBS and stained with directly conjugated antibodies for 30 minutes. Then the samples were rinsed with PBS 3X and mounted on cavity slides (Sigma) with Fluoromount G (eBioscience). The antibodies used are listed in **Table 3**. Approximately 50 µm thick Z-stack and tile scan were acquired with LSM880 Zeiss microscope with 40x/1.3 (oil). Images analysis was performed using Imaris (Bitplane) software. Adipocyte size was quantified using Bitplane Imaris image analysis software.

For perilipin staining, samples were collected in 4% methanol free PFA, fixed for 2 days at 4°C, and embedded in paraffin. 5 μm slides were cut with a Leica RM2265 microtome, dewax with xylene and rehydrated with ethanol bath of decreasing concentrations. Antigen retrieval was performed with pH 6 citrate solution (Cell Signaling). After endogenous peroxidase blocking for 10 min with 3% H_2_O_2_ solution (Sigma), the samples were blocked with TBS (Sigma) Tween 0.3% (Sigma), BSA 5%, Normal goat serum 5% (Sigma). Slides were incubated overnight at 4°C with anti-mouse and human perilipin antibody (1/200, clone D1D8, Cell Signaling). Secondary antibody, goat anti rabbit HRP coupled antibody (1/200, cell signaling) was incubated for 1 hour at room temperature before revelation with peroxidase substrate kit (Vector, SK4805), and counterstaining with hematoxylin (0.25%, Sigma). Slides were finally dehydrated in ethanol bath of increasing concentration and xylene baths and mounted in Entellan (Merck). Paraffin sections were also stained with Hematoxylin and eosin. Images were taken with different objectives (N-Achroplan 2.5x/0.07, N-Achroplan 10x/0.25, N-Achroplan 20x/0.45, N-Achroplan 63x/0.85) of a Axio Lab.A1 Zeiss. Gross morphology of the liver and adipose tissues were taken with a Leica M80. Crown-like structures (CLS) were quantified on the sections of H&E stained slides. For each sample, three to five regions were analyzed. Crown like structures were defined as microscopic foci of dying adipocytes surrounded by macrophages as visualized by H&E staining.

Analysis of macrophages was done on May-Grunwald Giemsa staining of sorted cells from each population. 1000 to 2000 cells for each sample were sorted directly in FBS (Invitrogen), and centrifuged for 10 min at 800 g with low acceleration onto Superfrost slides (Thermo Scientific). After air drying for at least 30 min, the slides were fixed in methanol, air dried for at least 30 min, stained with May-Grunwald (Sigma) solution for 5 to 15 min then Giemsa (Sigma) solution 14% for 15 to 30 min, and rinsed with Sorenson buffer pH 6.8. After air drying, the slides were mounted with Entellan (Merck). Pictures were taken using Axio Lab.A1 Zeiss microscope with N-Achroplan 100x/01.25 objective.

***In vitro* epididymal adipose tissue explants.** Epididymal fat pads of 4-day-old *Tnfrsf11a^Cre^; Csf1r^fl/fl^* mice and littermates were dissected and incubated in RPMI with 10% heat de-activated FBS at 37°c for 14 days(*18*). Fresh medium was added to the explants on day 7. At the termination of the experiment, the explants were fixed in 4% PFA for 30 minutes at room temperature with agitation and then rinsed in PBS and permeabilized for 15 minutes in PBS + 0.03% Triton. Subsequently, the explants were blocked with 2% BSA and stained with directly conjugated antibodies. To remove macrophages from *wt* epididymal fat pads *in vitro*, 15 μM of the CSF1R inhibitor, PLX5622, was added to explants on day 0 and 7 following the addition of fresh medium. For the *in* *vitro* rescue experiments, epididymal fat pads of *Tnfrsf11a^Cre^; Csf1r^fl/fl^* mice and littermates as well as PLX5266 treated fat pads were supplemented with 100 ng/ml of recombinant mouse PDGFcc (R&D systems), 10 ng/ml of recombinant mouse VEGFb (R&D systems), or 10 ng/ml of recombinant mouse IGF1 (R&D systems) three times on days 0, 3, and 6. Fluorescent while mount images were analyzed using Bitplabe Imaris image analysis software.

***Drosophila* lines and crosses.** Flies were raised at 25°C in a 12-h light/12-h dark cycle and maintained on food containing 10% w/v Brewer’s yeast, 8% fructose, 2% polenta and 0.8% Agar. Adult flies in vials were allowed to lay eggs for 2 hours, whereupon the adults were removed, and eggs were allowed to develop until the wandering larval stage.

| **Fly stock** | **Description** |
| --- | --- |
| *w^1118^; Srp^Hemo^Gal4* | Plasmatocyte specific line(*19*). A kind gift from Norbert Perrimon |
| *w^1118^;; uas-reaper* | A pro-apoptotic gene. BDSC # 50790 |
| *w^1118^;He-gal4* | BDSC #8699 |
| *w^1118^;uas-pvf3* | A kind gift from Michael Galko |
| *w^1118^;uas-dilp5-IR* | VDRC # 105004 |
| *w^1118^;uas-dpp-IR* | BDSC #25782 |
| *w^1118^;uas-pvf1-IR* | BDSC #39038 |
| *w^1118^;uas-pvf3-IR* | BDSC #38962 |
| *w^1118^;uas-upd3-IR* | VDRC # 106869 |
| *w^1118^;uas-Spaetzle-IR* | VDRC # 105017 |
| *y^1^ w^1118^* | BDSC #1495 |
| *y^1^ w^1118^;Pvf3^EY09531^* | BDSC #17577 |
| *w1118; srpHemo-3XmCherry* | A kind gift from Daria Siekhaus. |

**Triglyceride measurement, buoyancy assay, fat body cell size, and developmental timing in *Drosophila*.** To measure triglycerides, 10 wandering L3 larvae were homogenized in 100 μl of PBS + 0.05% Tween 20 with a pestle on ice. Triglyceride was then measured using the free glycerol reagent (Sigma; F6428) as described before(*20*) and normalized to the protein content. In a complementary experiment, the lipid content of the wandering L3 larvae were compared using buoyancy assay. The L3 larvae were placed in a 30% sucrose solution and then diluted slowly to a 8% sucrose solution with PBS. Afterwards, the larvae were left to equilibrate for 30 minutes at which point the position of the larvae in the solution was recorded.

To measure the size of the fat body cells, the larvae fat bodies were dissected out and then fixed in 4% PFA for 30 minutes at room temperature. The fixed tissues were then stained with Bodipy for 30 minutes and washed and imaged on an Zeiss LSM 880 with a 40X objective.

To examine the development of *Drosophila* larva, 30 eggs were seeded on freshly prepared food and then left to develop to the L3 stage.

**qPCR on sorted Drosophila hemocytes and whole larvae.** For qRT-PCR, 10000 hemocytes were sorted into 300μL of Trizol LS (Life Technologies). RNA extraction was performed following manufacturer’s instructions (Direct-zol RNA microPrep, Zymo Research). RNA concentration was measured with nanodrop2000. cDNA preparation was performed with Quantitect Reverse transcription kit (Qiagen) as per manufacturer’s instructions. qRT-PCR were done with 1ng cDNA.

For qRT-PCR performed on whole larvae, 5-10 larvae were homogenized in Trizol with a pestle on ice. RNA extraction and cDNA preparation was as described above. The same protocol as for the sorted cells and the qRT-PCR was performed on 10ng cDNA. qRT-PCR are performed on a Quant Studio 6 Flex using TaqMan Fast Advance Mastermix, and TaqMan probes for Gapdh2 (Dm01843776_s1), *dilp2* (Dm01822534_g1), *dilp4* (Dm01801938_g1), *dilp5* (Dm01798339_g1), *dilp6* (Dm01829746_g1), *Pvf1* (Dm01813949_m1), *Pvf3* (Dm01814376_m1), and *dpp* (Dm01842959_m1).

**Statistical tests.** Data are represented as individual values per mouse, unless otherwise stated. The *n* value represents biological replicates. Statistical significance was calculated using the software GraphPad Prism using unpaired Student's t-tests to compare 2 groups or one-way ANOVA, to compare more than 2 groups, as indicated in the figures. For RNA sequencing, R software was used, as described above.

**Data availability.** RNASeq data are available from GEO under the reference number GSE106084 (<https://www.ncbi.nlm.nih.gov/geo/query/acc.cgi?acc=GSE106084>). Other datasets that support the findings of this study are available from the corresponding author.
