## Supplementary material for "Diet-regulated production of PDGFcc by macrophages controls energy storage": Supp. Figures 1-9

### Slide 1
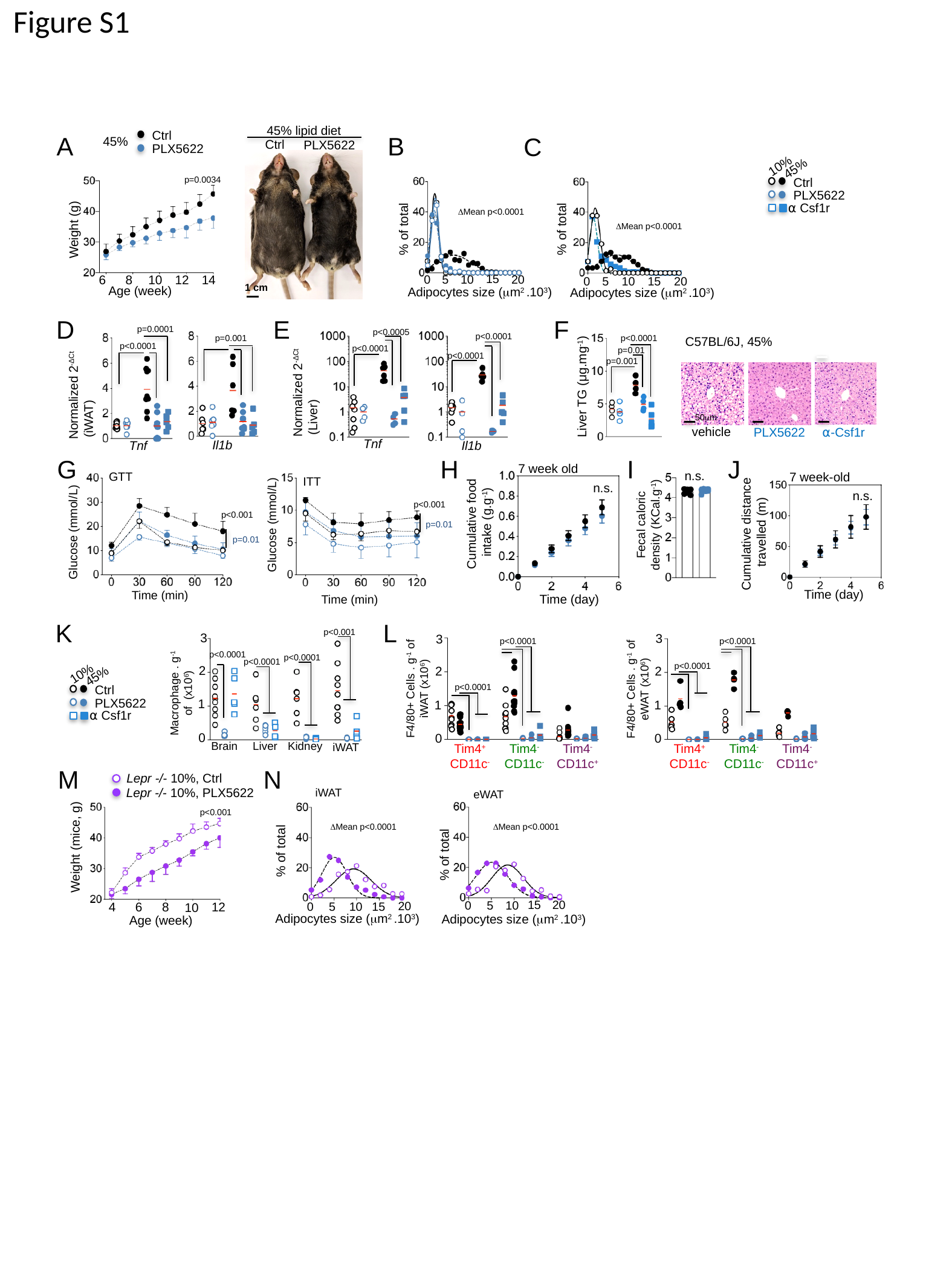

Figure S1
45% lipid diet
Ctrl
PLX5622
1 cm
Ctrl
45%
PLX5622
A
B
C
10%
45%
Ctrl
PLX5622
⍺ Csf1r
% of total
15
20
0
5
10
p=0.0034
Weight (g)
6
8
10
12
14
Age (week)
% of total
15
20
0
5
10
Adipocytes size (μm^2^ .10^3^)
ΔMean p<0.0001
ΔMean p<0.0001
Adipocytes size (μm^2^ .10^3^)
D
E
F
p=0.0001
p<0.0001
p<0.0001
p=0.01
p=0.001
Liver TG (μg.mg^-1^)
p<0.0005
p<0.0001
p=0.001
C57BL/6J, 45%
50μm
vehicle
⍺-Csf1r
PLX5622
p<0.0001
p<0.0001
Normalized 2^-ΔCt^
 (Liver)
Normalized 2^-ΔCt^
 (iWAT)
Tnf
Il1b
Il1b
Tnf
G
H
I
J
7 week old
n.s.
Cumulative food
intake (g.g^-1^)
Time (day)
n.s.
Fecal caloric density (KCal.g^-1^)
7 week-old
n.s.
Cumulative distance
 travelled (m)
Time (day)
GTT
ITT
p<0.001
p<0.001
p=0.01
Glucose (mmol/L)
Glucose (mmol/L)
p=0.01
Time (min)
Time (min)
K
L
3
p<0.0001
2
F4/80+ Cells . g^-1^ of iWAT (x10^6^)
p<0.0001
1
0
Tim4^+^
CD11c^-^
Tim4^-^ CD11c^-^
Tim4^-^ CD11c^+^
3
p<0.0001
p<0.0001
2
F4/80+ Cells . g^-1^ of eWAT (x10^6^)
1
0
Tim4^+^
CD11c^-^
Tim4^-^ CD11c^-^
Tim4^-^ CD11c^+^
p<0.001
3
p<0.0001
p<0.0001
p<0.0001
2
Macrophage . g^-1^ of (x10^6^)
1
0
Brain
Liver
Kidney
iWAT
10%
45%
Ctrl
PLX5622
⍺ Csf1r
M
N
 Lepr -/- 10%, Ctrl
Lepr -/- 10%, PLX5622
iWAT
eWAT
% of total
15
20
5
10
0
Weight (mice, g)
4
6
8
12
10
Age (week)
p<0.001
% of total
20
15
5
0
10
ΔMean p<0.0001
ΔMean p<0.0001
Adipocytes size (μm^2^ .10^3^)
Adipocytes size (μm^2^ .10^3^)

### Slide 2
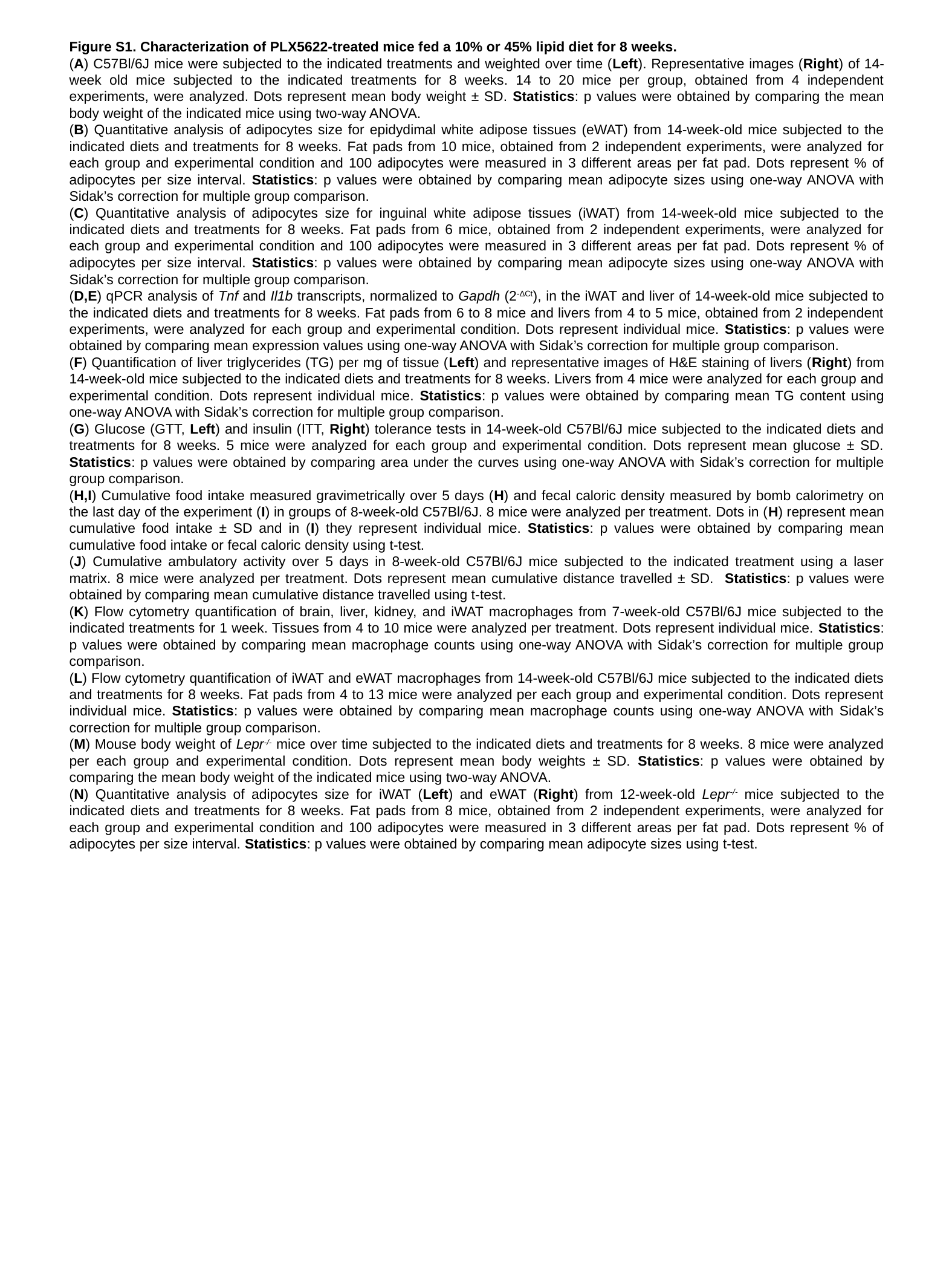

Figure S1. Characterization of PLX5622-treated mice fed a 10% or 45% lipid diet for 8 weeks.
(A) C57Bl/6J mice were subjected to the indicated treatments and weighted over time (Left). Representative images (Right) of 14-week old mice subjected to the indicated treatments for 8 weeks. 14 to 20 mice per group, obtained from 4 independent experiments, were analyzed. Dots represent mean body weight ± SD. Statistics: p values were obtained by comparing the mean body weight of the indicated mice using two-way ANOVA.
(B) Quantitative analysis of adipocytes size for epidydimal white adipose tissues (eWAT) from 14-week-old mice subjected to the indicated diets and treatments for 8 weeks. Fat pads from 10 mice, obtained from 2 independent experiments, were analyzed for each group and experimental condition and 100 adipocytes were measured in 3 different areas per fat pad. Dots represent % of adipocytes per size interval. Statistics: p values were obtained by comparing mean adipocyte sizes using one-way ANOVA with Sidak’s correction for multiple group comparison.
(C) Quantitative analysis of adipocytes size for inguinal white adipose tissues (iWAT) from 14-week-old mice subjected to the indicated diets and treatments for 8 weeks. Fat pads from 6 mice, obtained from 2 independent experiments, were analyzed for each group and experimental condition and 100 adipocytes were measured in 3 different areas per fat pad. Dots represent % of adipocytes per size interval. Statistics: p values were obtained by comparing mean adipocyte sizes using one-way ANOVA with Sidak’s correction for multiple group comparison.
(D,E) qPCR analysis of Tnf and Il1b transcripts, normalized to Gapdh (2^-ΔCt^), in the iWAT and liver of 14-week-old mice subjected to the indicated diets and treatments for 8 weeks. Fat pads from 6 to 8 mice and livers from 4 to 5 mice, obtained from 2 independent experiments, were analyzed for each group and experimental condition. Dots represent individual mice. Statistics: p values were obtained by comparing mean expression values using one-way ANOVA with Sidak’s correction for multiple group comparison.
(F) Quantification of liver triglycerides (TG) per mg of tissue (Left) and representative images of H&E staining of livers (Right) from 14-week-old mice subjected to the indicated diets and treatments for 8 weeks. Livers from 4 mice were analyzed for each group and experimental condition. Dots represent individual mice. Statistics: p values were obtained by comparing mean TG content using one-way ANOVA with Sidak’s correction for multiple group comparison.
(G) Glucose (GTT, Left) and insulin (ITT, Right) tolerance tests in 14-week-old C57Bl/6J mice subjected to the indicated diets and treatments for 8 weeks. 5 mice were analyzed for each group and experimental condition. Dots represent mean glucose ± SD. Statistics: p values were obtained by comparing area under the curves using one-way ANOVA with Sidak’s correction for multiple group comparison.
(H,I) Cumulative food intake measured gravimetrically over 5 days (H) and fecal caloric density measured by bomb calorimetry on the last day of the experiment (I) in groups of 8-week-old C57Bl/6J. 8 mice were analyzed per treatment. Dots in (H) represent mean cumulative food intake ± SD and in (I) they represent individual mice. Statistics: p values were obtained by comparing mean cumulative food intake or fecal caloric density using t-test.
(J) Cumulative ambulatory activity over 5 days in 8-week-old C57Bl/6J mice subjected to the indicated treatment using a laser matrix. 8 mice were analyzed per treatment. Dots represent mean cumulative distance travelled ± SD. Statistics: p values were obtained by comparing mean cumulative distance travelled using t-test.
(K) Flow cytometry quantification of brain, liver, kidney, and iWAT macrophages from 7-week-old C57Bl/6J mice subjected to the indicated treatments for 1 week. Tissues from 4 to 10 mice were analyzed per treatment. Dots represent individual mice. Statistics: p values were obtained by comparing mean macrophage counts using one-way ANOVA with Sidak’s correction for multiple group comparison.
(L) Flow cytometry quantification of iWAT and eWAT macrophages from 14-week-old C57Bl/6J mice subjected to the indicated diets and treatments for 8 weeks. Fat pads from 4 to 13 mice were analyzed per each group and experimental condition. Dots represent individual mice. Statistics: p values were obtained by comparing mean macrophage counts using one-way ANOVA with Sidak’s correction for multiple group comparison.
(M) Mouse body weight of Lepr^-/-^ mice over time subjected to the indicated diets and treatments for 8 weeks. 8 mice were analyzed per each group and experimental condition. Dots represent mean body weights ± SD. Statistics: p values were obtained by comparing the mean body weight of the indicated mice using two-way ANOVA.
(N) Quantitative analysis of adipocytes size for iWAT (Left) and eWAT (Right) from 12-week-old Lepr^-/-^ mice subjected to the indicated diets and treatments for 8 weeks. Fat pads from 8 mice, obtained from 2 independent experiments, were analyzed for each group and experimental condition and 100 adipocytes were measured in 3 different areas per fat pad. Dots represent % of adipocytes per size interval. Statistics: p values were obtained by comparing mean adipocyte sizes using t-test.

### Slide 3
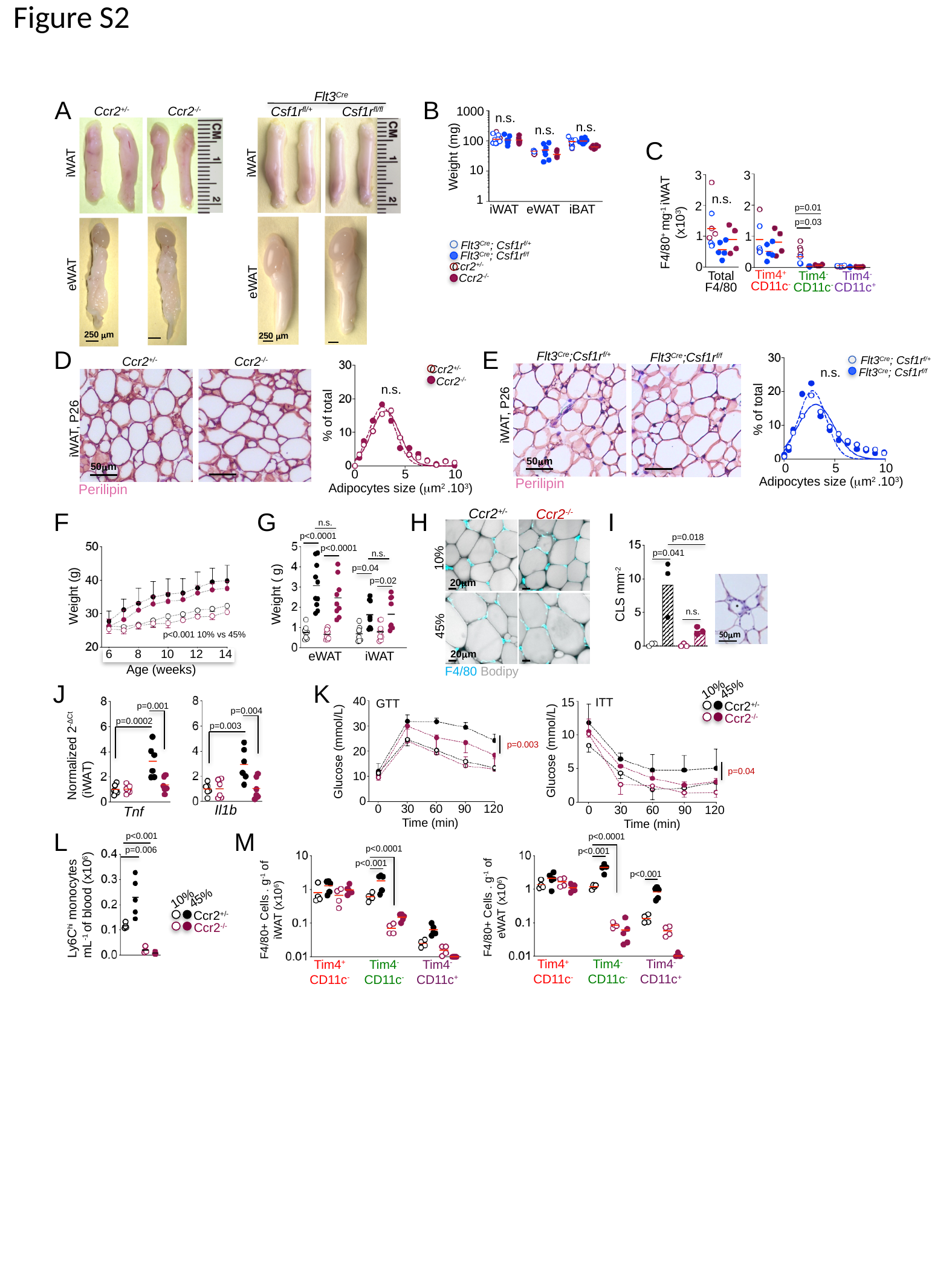

Figure S2
Flt3^Cre^
A
B
1000
100
Weight (mg)
10
1
iWAT
eWAT
iBAT
Csf1r^fl/+^
Csf1r^fl/fl^
Ccr2^+/-^
Ccr2^-/-^
n.s.
n.s.
n.s.
C
iWAT
iWAT
3
3
2
2
p=0.01
F4/80^+^ mg^-1^ iWAT (x10^3^)
p=0.03
1
1
0
0
Total F4/80
Tim4^-^ CD11c^-^
Tim4^-^ CD11c^+^
Tim4^+^
CD11c^-^
n.s.
250 μm
Flt3^Cre^; Csf1r^f/+^
Flt3^Cre^; Csf1r^f/f^
Ccr2^+/-^
Ccr2^-/-^
eWAT
eWAT
250 μm
D
E
Flt3^Cre^;Csf1r^f/+^
Flt3^Cre^;Csf1r^f/f^
iWAT, P26
 50μm
% of total
0
5
10
Adipocytes size (μm^2^ .10^3^)
Flt3^Cre^; Csf1r^f/+^
Flt3^Cre^; Csf1r^f/f^
Ccr2^+/-^
Ccr2^-/-^
iWAT, P26
Perilipin
 50μm
% of total
0
5
10
Adipocytes size (μm^2^ .10^3^)
Ccr2^+/-^
Ccr2^-/-^
n.s.
n.s.
Perilipin
Ccr2^+/-^
Ccr2^-/-^
10%
20μm
45%
20μm
F
G
H
I
n.s.
p<0.0001
p<0.0001
n.s.
p=0.04
p=0.02
Weight ( g)
eWAT
iWAT
p=0.018
Weight (g)
p<0.001 10% vs 45%
6
8
10
12
14
Age (weeks)
p=0.041
CLS mm^-2^
n.s.
*
50μm
F4/80 Bodipy
J
K
45%
10%
Ccr2^+/-^
Ccr2^-/-^
p=0.004
p=0.003
Il1b
p=0.001
p=0.0002
Normalized 2^-ΔCt^
 (iWAT)
Tnf
40
GTT
30
p=0.003
20
Glucose (mmol/L)
10
0
120
90
60
30
0
Time (min)
15
ITT
10
5
p=0.04
0
120
90
60
30
0
Time (min)
Glucose (mmol/L)
L
M
p<0.001
p<0.0001
p<0.001
p<0.001
Tim4^+^
CD11c^-^
Tim4^-^ CD11c^-^
Tim4^-^ CD11c^+^
p<0.0001
p<0.001
Tim4^+^
CD11c^-^
Tim4^-^ CD11c^-^
Tim4^-^ CD11c^+^
p=0.006
Ly6C^hi^ monocytes
mL^-1^ of blood (x10^6^)
45%
10%
Ccr2^+/-^
Ccr2^-/-^
F4/80+ Cells . g^-1^ of eWAT (x10^6^)
F4/80+ Cells . g^-1^ of iWAT (x10^6^)

### Slide 4
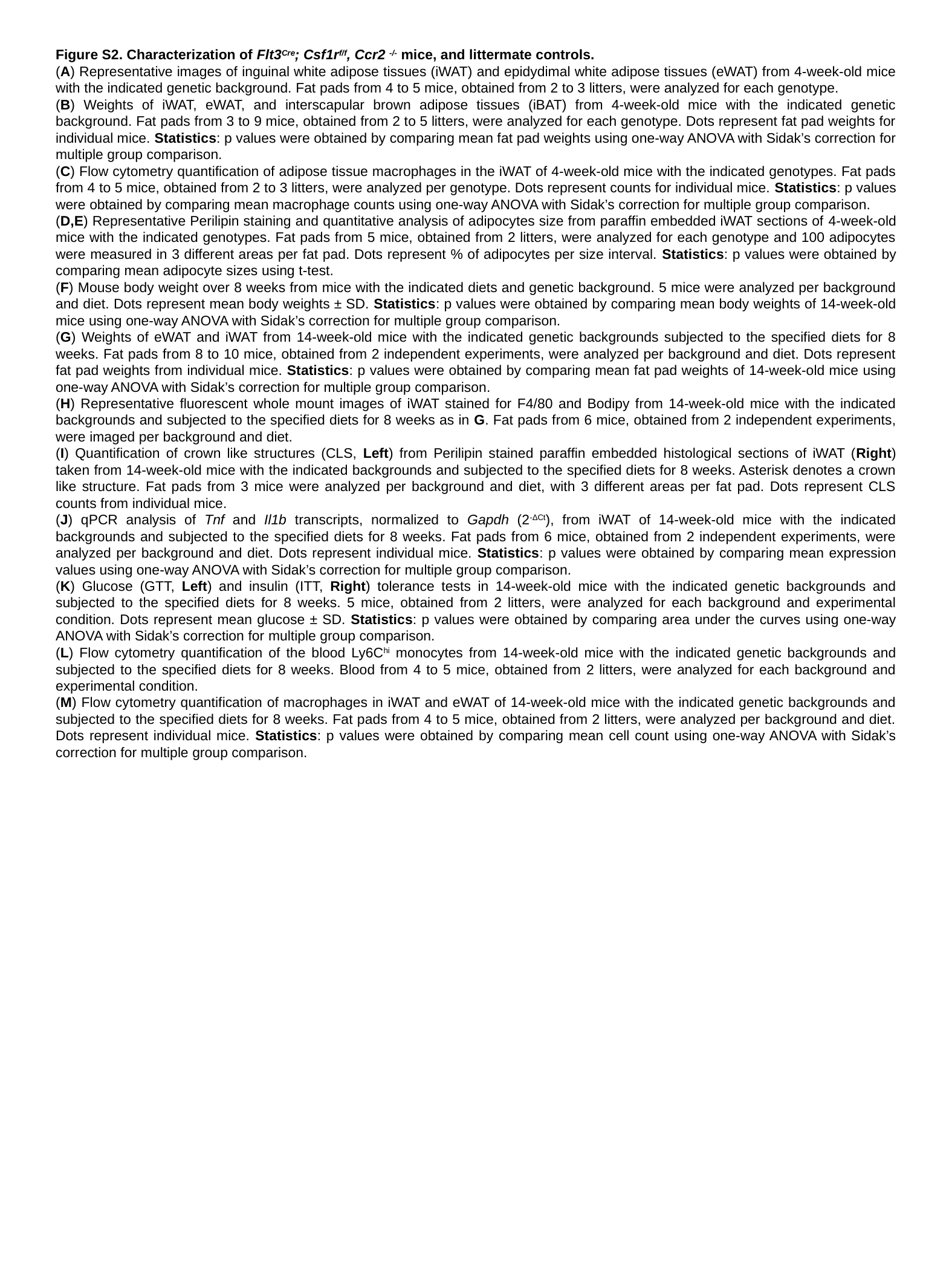

Figure S2. Characterization of Flt3^Cre^; Csf1r^f/f^, Ccr2 ^-/-^ mice, and littermate controls.
(A) Representative images of inguinal white adipose tissues (iWAT) and epidydimal white adipose tissues (eWAT) from 4-week-old mice with the indicated genetic background. Fat pads from 4 to 5 mice, obtained from 2 to 3 litters, were analyzed for each genotype.
(B) Weights of iWAT, eWAT, and interscapular brown adipose tissues (iBAT) from 4-week-old mice with the indicated genetic background. Fat pads from 3 to 9 mice, obtained from 2 to 5 litters, were analyzed for each genotype. Dots represent fat pad weights for individual mice. Statistics: p values were obtained by comparing mean fat pad weights using one-way ANOVA with Sidak’s correction for multiple group comparison.
(C) Flow cytometry quantification of adipose tissue macrophages in the iWAT of 4-week-old mice with the indicated genotypes. Fat pads from 4 to 5 mice, obtained from 2 to 3 litters, were analyzed per genotype. Dots represent counts for individual mice. Statistics: p values were obtained by comparing mean macrophage counts using one-way ANOVA with Sidak’s correction for multiple group comparison.
(D,E) Representative Perilipin staining and quantitative analysis of adipocytes size from paraffin embedded iWAT sections of 4-week-old mice with the indicated genotypes. Fat pads from 5 mice, obtained from 2 litters, were analyzed for each genotype and 100 adipocytes were measured in 3 different areas per fat pad. Dots represent % of adipocytes per size interval. Statistics: p values were obtained by comparing mean adipocyte sizes using t-test.
(F) Mouse body weight over 8 weeks from mice with the indicated diets and genetic background. 5 mice were analyzed per background and diet. Dots represent mean body weights ± SD. Statistics: p values were obtained by comparing mean body weights of 14-week-old mice using one-way ANOVA with Sidak’s correction for multiple group comparison.
(G) Weights of eWAT and iWAT from 14-week-old mice with the indicated genetic backgrounds subjected to the specified diets for 8 weeks. Fat pads from 8 to 10 mice, obtained from 2 independent experiments, were analyzed per background and diet. Dots represent fat pad weights from individual mice. Statistics: p values were obtained by comparing mean fat pad weights of 14-week-old mice using one-way ANOVA with Sidak’s correction for multiple group comparison.
(H) Representative fluorescent whole mount images of iWAT stained for F4/80 and Bodipy from 14-week-old mice with the indicated backgrounds and subjected to the specified diets for 8 weeks as in G. Fat pads from 6 mice, obtained from 2 independent experiments, were imaged per background and diet.
(I) Quantification of crown like structures (CLS, Left) from Perilipin stained paraffin embedded histological sections of iWAT (Right) taken from 14-week-old mice with the indicated backgrounds and subjected to the specified diets for 8 weeks. Asterisk denotes a crown like structure. Fat pads from 3 mice were analyzed per background and diet, with 3 different areas per fat pad. Dots represent CLS counts from individual mice.
(J) qPCR analysis of Tnf and Il1b transcripts, normalized to Gapdh (2^-ΔCt^), from iWAT of 14-week-old mice with the indicated backgrounds and subjected to the specified diets for 8 weeks. Fat pads from 6 mice, obtained from 2 independent experiments, were analyzed per background and diet. Dots represent individual mice. Statistics: p values were obtained by comparing mean expression values using one-way ANOVA with Sidak’s correction for multiple group comparison.
(K) Glucose (GTT, Left) and insulin (ITT, Right) tolerance tests in 14-week-old mice with the indicated genetic backgrounds and subjected to the specified diets for 8 weeks. 5 mice, obtained from 2 litters, were analyzed for each background and experimental condition. Dots represent mean glucose ± SD. Statistics: p values were obtained by comparing area under the curves using one-way ANOVA with Sidak’s correction for multiple group comparison.
(L) Flow cytometry quantification of the blood Ly6C^hi^ monocytes from 14-week-old mice with the indicated genetic backgrounds and subjected to the specified diets for 8 weeks. Blood from 4 to 5 mice, obtained from 2 litters, were analyzed for each background and experimental condition.
(M) Flow cytometry quantification of macrophages in iWAT and eWAT of 14-week-old mice with the indicated genetic backgrounds and subjected to the specified diets for 8 weeks. Fat pads from 4 to 5 mice, obtained from 2 litters, were analyzed per background and diet. Dots represent individual mice. Statistics: p values were obtained by comparing mean cell count using one-way ANOVA with Sidak’s correction for multiple group comparison.

### Slide 5
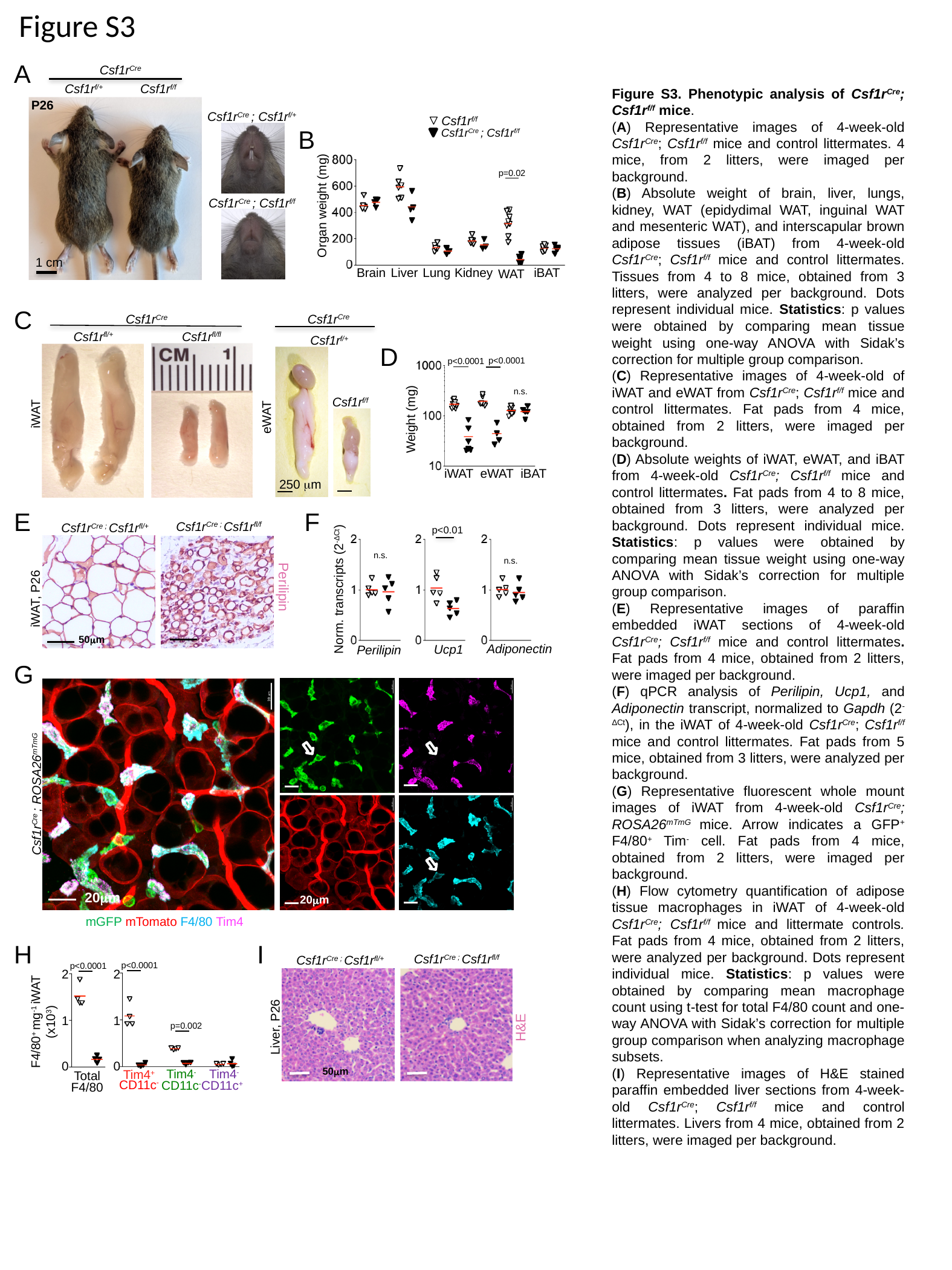

Figure S3
A
Csf1r^Cre^
Csf1r^f/+^
Csf1r^f/f^
P26
eWAT
1 cm
Figure S3. Phenotypic analysis of Csf1r^Cre^; Csf1r^f/f^ mice.
(A) Representative images of 4-week-old Csf1r^Cre^; Csf1r^f/f^ mice and control littermates. 4 mice, from 2 litters, were imaged per background.
(B) Absolute weight of brain, liver, lungs, kidney, WAT (epidydimal WAT, inguinal WAT and mesenteric WAT), and interscapular brown adipose tissues (iBAT) from 4-week-old Csf1r^Cre^; Csf1r^f/f^ mice and control littermates. Tissues from 4 to 8 mice, obtained from 3 litters, were analyzed per background. Dots represent individual mice. Statistics: p values were obtained by comparing mean tissue weight using one-way ANOVA with Sidak’s correction for multiple group comparison.
(C) Representative images of 4-week-old of iWAT and eWAT from Csf1r^Cre^; Csf1r^f/f^ mice and control littermates. Fat pads from 4 mice, obtained from 2 litters, were imaged per background.
(D) Absolute weights of iWAT, eWAT, and iBAT from 4-week-old Csf1r^Cre^; Csf1r^f/f^ mice and control littermates. Fat pads from 4 to 8 mice, obtained from 3 litters, were analyzed per background. Dots represent individual mice. Statistics: p values were obtained by comparing mean tissue weight using one-way ANOVA with Sidak’s correction for multiple group comparison.
(E) Representative images of paraffin embedded iWAT sections of 4-week-old Csf1r^Cre^; Csf1r^f/f^ mice and control littermates. Fat pads from 4 mice, obtained from 2 litters, were imaged per background.
(F) qPCR analysis of Perilipin, Ucp1, and Adiponectin transcript, normalized to Gapdh (2^-ΔCt^), in the iWAT of 4-week-old Csf1r^Cre^; Csf1r^f/f^ mice and control littermates. Fat pads from 5 mice, obtained from 3 litters, were analyzed per background.
(G) Representative fluorescent whole mount images of iWAT from 4-week-old Csf1r^Cre^; ROSA26^mTmG^ mice. Arrow indicates a GFP^+^ F4/80^+^ Tim^-^ cell. Fat pads from 4 mice, obtained from 2 litters, were imaged per background.
(H) Flow cytometry quantification of adipose tissue macrophages in iWAT of 4-week-old Csf1r^Cre^; Csf1r^f/f^ mice and littermate controls. Fat pads from 4 mice, obtained from 2 litters, were analyzed per background. Dots represent individual mice. Statistics: p values were obtained by comparing mean macrophage count using t-test for total F4/80 count and one-way ANOVA with Sidak’s correction for multiple group comparison when analyzing macrophage subsets.
(I) Representative images of H&E stained paraffin embedded liver sections from 4-week-old Csf1r^Cre^; Csf1r^f/f^ mice and control littermates. Livers from 4 mice, obtained from 2 litters, were imaged per background.
Csf1r^Cre^ ; Csf1r^f/+^
Csf1r^f/f^
Csf1r^Cre^ ; Csf1r^f/f^
B
p=0.02
Organ weight (mg)
Brain
Liver
Lung
Kidney
iBAT
WAT
Csf1r^Cre^ ; Csf1r^f/f^
C
Csf1r^Cre^
Csf1r^f/+^
Csf1r^f/f^
250 μm
eWAT
Csf1r^Cre^
Csf1r^fl/+^
Csf1r^fl/fl^
iWAT
D
p<0.0001
p<0.0001
Weight (mg)
iWAT
eWAT
iBAT
n.s.
E
F
Csf1r^Cre ;^ Csf1r^fl/f^
Csf1r^Cre ;^ Csf1r^fl/+^
p<0.01
n.s.
n.s.
Norm. transcripts (2^-ΔCt^)
iWAT, P26
Perilipin
Csf1r^f/f^
Csf1r^Cre+^ ; Csf1r^f/f^
 50μm
Adiponectin
Ucp1
Perilipin
G
Csf1r^Cre^ ; ROSA26^mTmG^
20μm
20μm
mGFP mTomato F4/80 Tim4
H
I
Csf1r^Cre ;^ Csf1r^fl/f^
Csf1r^Cre ;^ Csf1r^fl/+^
p<0.0001
p<0.0001
2
1
0
2
1
0
F4/80^+^ mg^-1^ iWAT (x10^3^)
p=0.002
Tim4^-^ CD11c^-^
Tim4^-^ CD11c^+^
Tim4^+^
CD11c^-^
Total F4/80
H&E
Liver, P26
 50μm

### Slide 6
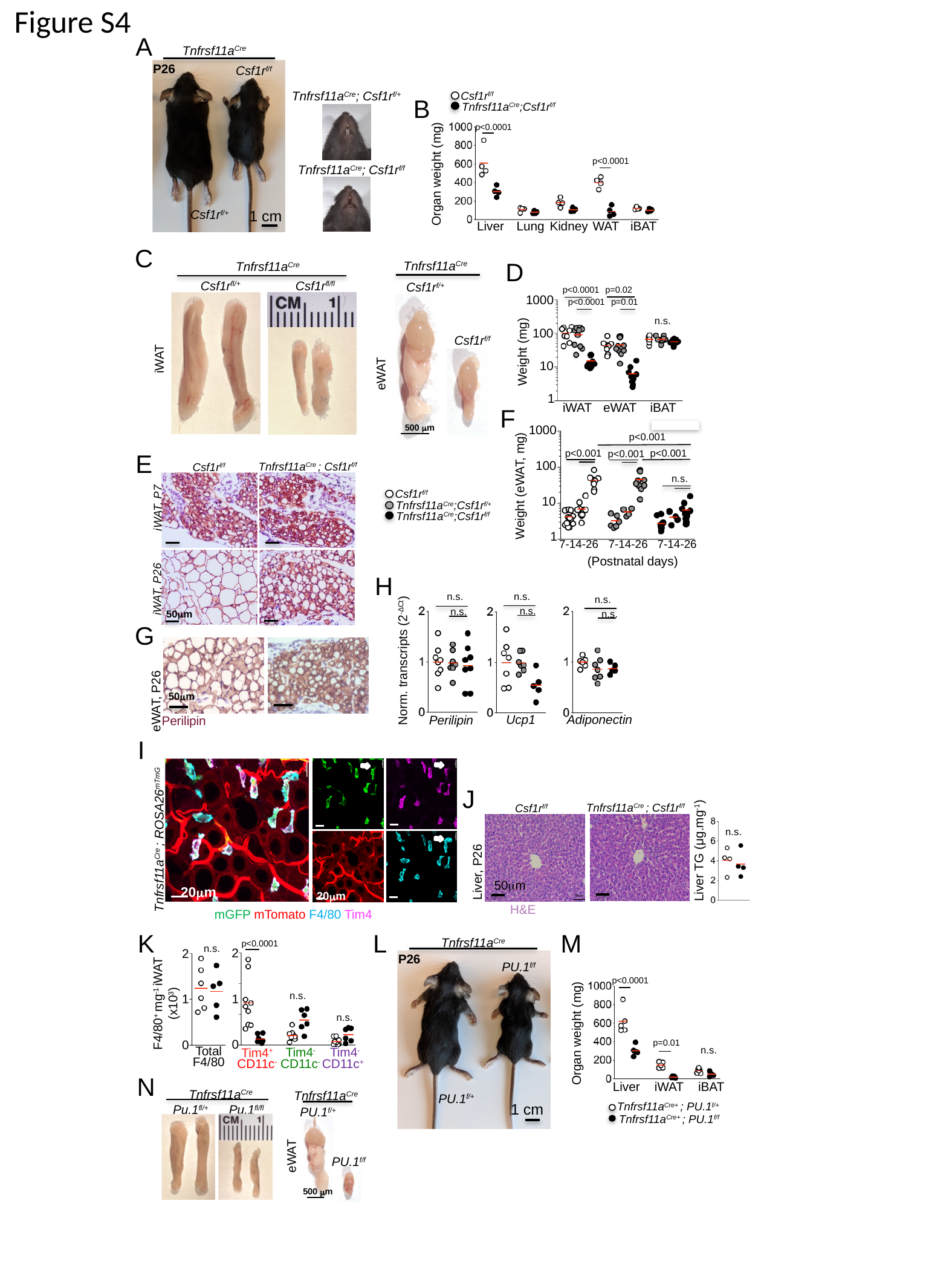

I
G
Figure S4
A
Tnfrsf11a^Cre^
P26
Csf1r^f/f^
1 cm
Csf1r^f/+^
p<0.0001
p<0.0001
Organ weight (mg)
Liver
Lung
Kidney
WAT
iBAT
Tnfrsf11a^Cre^; Csf1r^f/+^
Csf1r^f/f^
Tnfrsf11a^Cre^;Csf1r^f/f^
B
Tnfrsf11a^Cre^; Csf1r^f/f^
C
D
Tnfrsf11a^Cre^
Csf1r^f/+^
Csf1r^f/f^
500 μm
eWAT
Tnfrsf11a^Cre^
Csf1r^fl/+^
Csf1r^fl/fl^
iWAT
p<0.0001
p=0.02
1000
p<0.0001
p=0.01
100
Weight (mg)
10
1
eWAT
iBAT
iWAT
n.s.
F
1000
p<0.001
p<0.001
p<0.001
p<0.001
100
n.s.
Weight (eWAT, mg)
10
1
7-14-26
7-14-26
7-14-26
(Postnatal days)
E
Tnfrsf11a^Cre^ ; Csf1r^f/f^
Csf1r^f/f^
Csf1r^f/f^
Tnfrsf11a^Cre^;Csf1r^f/+^
Tnfrsf11a^Cre^;Csf1r^f/f^
iWAT, P7
H
iWAT, P26
n.s.
n.s.
n.s.
n.s.
n.s.
50μm
n.s.
G
Norm. transcripts (2^-ΔCt^)
50μm
eWAT, P26
Ucp1
Adiponectin
Perilipin
Perilipin
I
J
Liver TG (μg.mg^-1^)
Tnfrsf11a^Cre^ ; Csf1r^f/f^
Csf1r^f/f^
n.s.
Tnfrsf11a^Cre^ ; ROSA26^mTmG^
Liver, P26
50μm
20μm
20μm
H&E
mGFP mTomato F4/80 Tim4
K
L
M
Tnfrsf11a^Cre^
P26
PU.1^f/f^
PU.1^f/+^
1 cm
p<0.0001
2
2
1
0
F4/80^+^ mg^-1^ iWAT (x10^3^)
1
0
Total F4/80
Tim4^-^ CD11c^-^
Tim4^-^ CD11c^+^
Tim4^+^
CD11c^-^
n.s.
p<0.0001
p=0.01
Liver
iWAT
iBAT
Organ weight (mg)
n.s.
n.s.
n.s.
N
Tnfrsf11a^Cre^
Pu.1^fl/fl^
Pu.1^fl/+^
Tnfrsf11a^Cre^
PU.1^f/+^
PU.1^f/f^
500 μm
eWAT
Tnfrsf11a^Cre+^ ; PU.1^f/+^
Tnfrsf11a^Cre+^ ; PU.1^f/f^
L

### Slide 7
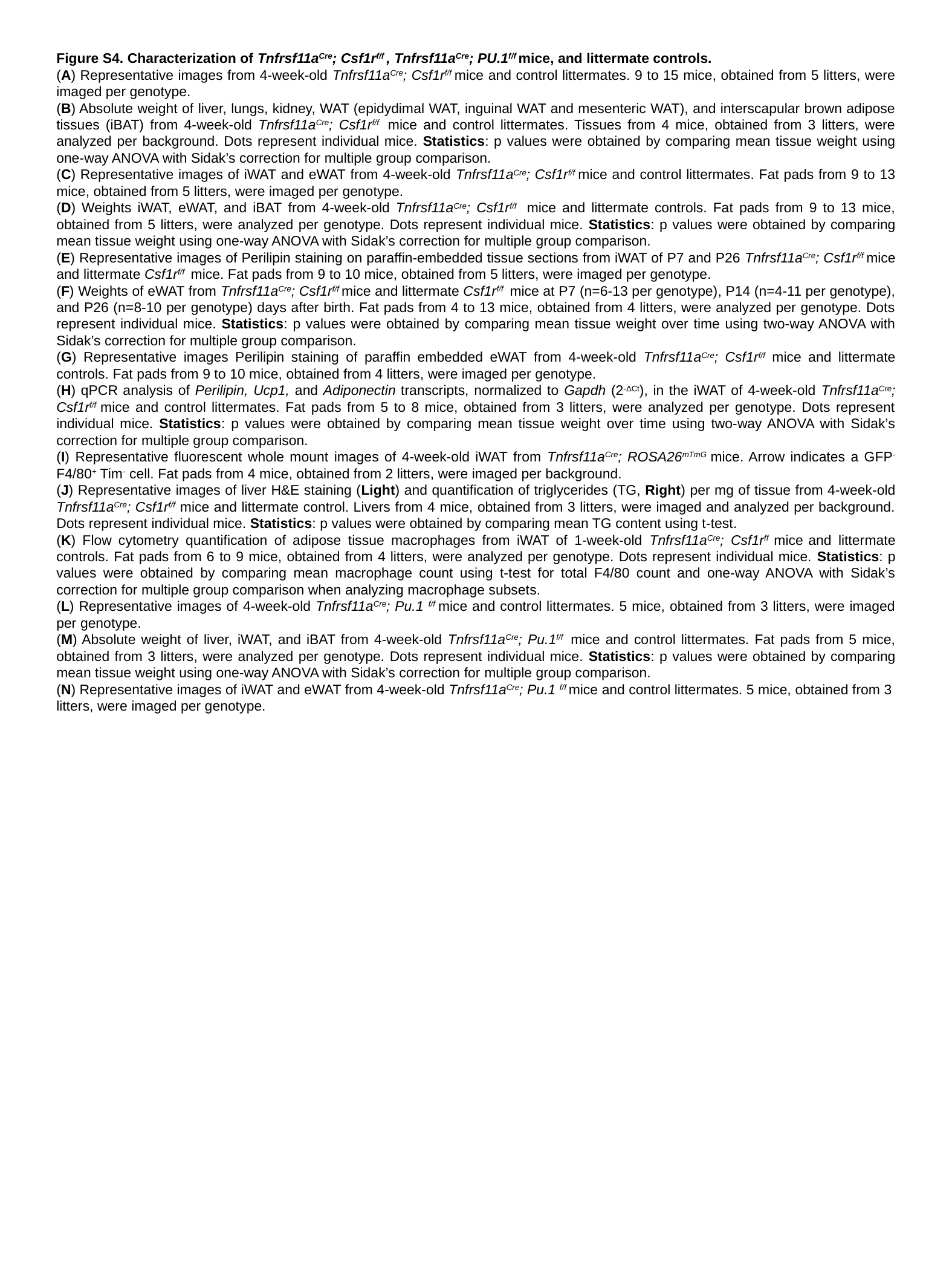

Figure S4. Characterization of Tnfrsf11a^Cre^; Csf1r^f/f^ , Tnfrsf11a^Cre^; PU.1^f/f^ mice, and littermate controls.
(A) Representative images from 4-week-old Tnfrsf11a^Cre^; Csf1r^f/f^ mice and control littermates. 9 to 15 mice, obtained from 5 litters, were imaged per genotype.
(B) Absolute weight of liver, lungs, kidney, WAT (epidydimal WAT, inguinal WAT and mesenteric WAT), and interscapular brown adipose tissues (iBAT) from 4-week-old Tnfrsf11a^Cre^; Csf1r^f/f^ mice and control littermates. Tissues from 4 mice, obtained from 3 litters, were analyzed per background. Dots represent individual mice. Statistics: p values were obtained by comparing mean tissue weight using one-way ANOVA with Sidak’s correction for multiple group comparison.
(C) Representative images of iWAT and eWAT from 4-week-old Tnfrsf11a^Cre^; Csf1r^f/f^ mice and control littermates. Fat pads from 9 to 13 mice, obtained from 5 litters, were imaged per genotype.
(D) Weights iWAT, eWAT, and iBAT from 4-week-old Tnfrsf11a^Cre^; Csf1r^f/f^ mice and littermate controls. Fat pads from 9 to 13 mice, obtained from 5 litters, were analyzed per genotype. Dots represent individual mice. Statistics: p values were obtained by comparing mean tissue weight using one-way ANOVA with Sidak’s correction for multiple group comparison.
(E) Representative images of Perilipin staining on paraffin-embedded tissue sections from iWAT of P7 and P26 Tnfrsf11a^Cre^; Csf1r^f/f^ mice and littermate Csf1r^f/f^ mice. Fat pads from 9 to 10 mice, obtained from 5 litters, were imaged per genotype.
(F) Weights of eWAT from Tnfrsf11a^Cre^; Csf1r^f/f^ mice and littermate Csf1r^f/f^ mice at P7 (n=6-13 per genotype), P14 (n=4-11 per genotype), and P26 (n=8-10 per genotype) days after birth. Fat pads from 4 to 13 mice, obtained from 4 litters, were analyzed per genotype. Dots represent individual mice. Statistics: p values were obtained by comparing mean tissue weight over time using two-way ANOVA with Sidak’s correction for multiple group comparison.
(G) Representative images Perilipin staining of paraffin embedded eWAT from 4-week-old Tnfrsf11a^Cre^; Csf1r^f/f^ mice and littermate controls. Fat pads from 9 to 10 mice, obtained from 4 litters, were imaged per genotype.
(H) qPCR analysis of Perilipin, Ucp1, and Adiponectin transcripts, normalized to Gapdh (2^-ΔCt^), in the iWAT of 4-week-old Tnfrsf11a^Cre^; Csf1r^f/f^ mice and control littermates. Fat pads from 5 to 8 mice, obtained from 3 litters, were analyzed per genotype. Dots represent individual mice. Statistics: p values were obtained by comparing mean tissue weight over time using two-way ANOVA with Sidak’s correction for multiple group comparison.
(I) Representative fluorescent whole mount images of 4-week-old iWAT from Tnfrsf11a^Cre^; ROSA26^mTmG^ mice. Arrow indicates a GFP^-^ F4/80^+^ Tim^-^ cell. Fat pads from 4 mice, obtained from 2 litters, were imaged per background.
(J) Representative images of liver H&E staining (Light) and quantification of triglycerides (TG, Right) per mg of tissue from 4-week-old Tnfrsf11a^Cre^; Csf1r^f/f^ mice and littermate control. Livers from 4 mice, obtained from 3 litters, were imaged and analyzed per background. Dots represent individual mice. Statistics: p values were obtained by comparing mean TG content using t-test.
(K) Flow cytometry quantification of adipose tissue macrophages from iWAT of 1-week-old Tnfrsf11a^Cre^; Csf1r^ff^ mice and littermate controls. Fat pads from 6 to 9 mice, obtained from 4 litters, were analyzed per genotype. Dots represent individual mice. Statistics: p values were obtained by comparing mean macrophage count using t-test for total F4/80 count and one-way ANOVA with Sidak’s correction for multiple group comparison when analyzing macrophage subsets.
(L) Representative images of 4-week-old Tnfrsf11a^Cre^; Pu.1 ^f/f^ mice and control littermates. 5 mice, obtained from 3 litters, were imaged per genotype.
(M) Absolute weight of liver, iWAT, and iBAT from 4-week-old Tnfrsf11a^Cre^; Pu.1^f/f^ mice and control littermates. Fat pads from 5 mice, obtained from 3 litters, were analyzed per genotype. Dots represent individual mice. Statistics: p values were obtained by comparing mean tissue weight using one-way ANOVA with Sidak’s correction for multiple group comparison.
(N) Representative images of iWAT and eWAT from 4-week-old Tnfrsf11a^Cre^; Pu.1 ^f/f^ mice and control littermates. 5 mice, obtained from 3 litters, were imaged per genotype.

### Slide 8
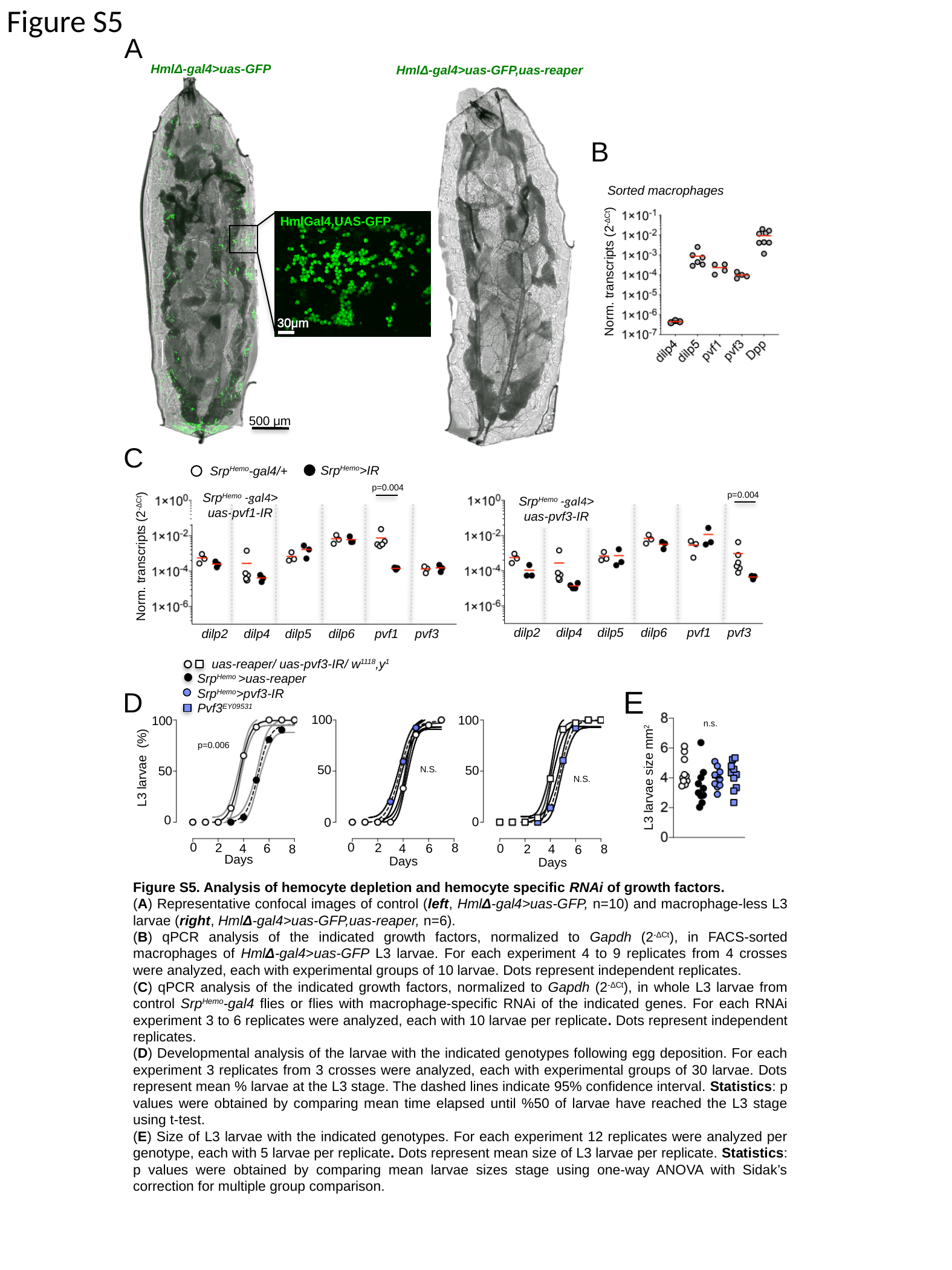

Figure S5
A
HmlΔ-gal4>uas-GFP
HmlΔ-gal4>uas-GFP,uas-reaper
B
Sorted macrophages
Norm. transcripts (2^-ΔCt^)
HmlGal4,UAS-GFP
 30μm
 500 μm
C
Srp^Hemo^>IR
Srp^Hemo^-gal4/+
p=0.004
p=0.004
Srp^Hemo^ -gal4>
uas-pvf1-IR
Srp^Hemo^ -gal4>
uas-pvf3-IR
Norm. transcripts (2^-ΔCt^)
dilp2
dilp4
dilp5
dilp6
pvf1
pvf3
dilp2
dilp4
dilp5
dilp6
pvf1
pvf3
uas-reaper/ uas-pvf3-IR/ w^1118^,y^1^
Srp^Hemo^ >uas-reaper
Pvf3^EY09531^
Srp^Hemo^>pvf3-IR
E
D
100
50
0
0
8
2
4
6
Days
100
50
0
2
4
6
8
Days
100
50
0
0
8
2
4
6
Days
n.s.
p=0.006
L3 larvae (%)
N.S.
L3 larvae size mm^2^
N.S.
0
Figure S5. Analysis of hemocyte depletion and hemocyte specific RNAi of growth factors.
(A) Representative confocal images of control (left, HmlΔ-gal4>uas-GFP, n=10) and macrophage-less L3 larvae (right, HmlΔ-gal4>uas-GFP,uas-reaper, n=6).
(B) qPCR analysis of the indicated growth factors, normalized to Gapdh (2^-ΔCt^), in FACS-sorted macrophages of HmlΔ-gal4>uas-GFP L3 larvae. For each experiment 4 to 9 replicates from 4 crosses were analyzed, each with experimental groups of 10 larvae. Dots represent independent replicates.
(C) qPCR analysis of the indicated growth factors, normalized to Gapdh (2^-ΔCt^), in whole L3 larvae from control Srp^Hemo^-gal4 flies or flies with macrophage-specific RNAi of the indicated genes. For each RNAi experiment 3 to 6 replicates were analyzed, each with 10 larvae per replicate. Dots represent independent replicates.
(D) Developmental analysis of the larvae with the indicated genotypes following egg deposition. For each experiment 3 replicates from 3 crosses were analyzed, each with experimental groups of 30 larvae. Dots represent mean % larvae at the L3 stage. The dashed lines indicate 95% confidence interval. Statistics: p values were obtained by comparing mean time elapsed until %50 of larvae have reached the L3 stage using t-test.
(E) Size of L3 larvae with the indicated genotypes. For each experiment 12 replicates were analyzed per genotype, each with 5 larvae per replicate. Dots represent mean size of L3 larvae per replicate. Statistics: p values were obtained by comparing mean larvae sizes stage using one-way ANOVA with Sidak’s correction for multiple group comparison.

### Slide 9
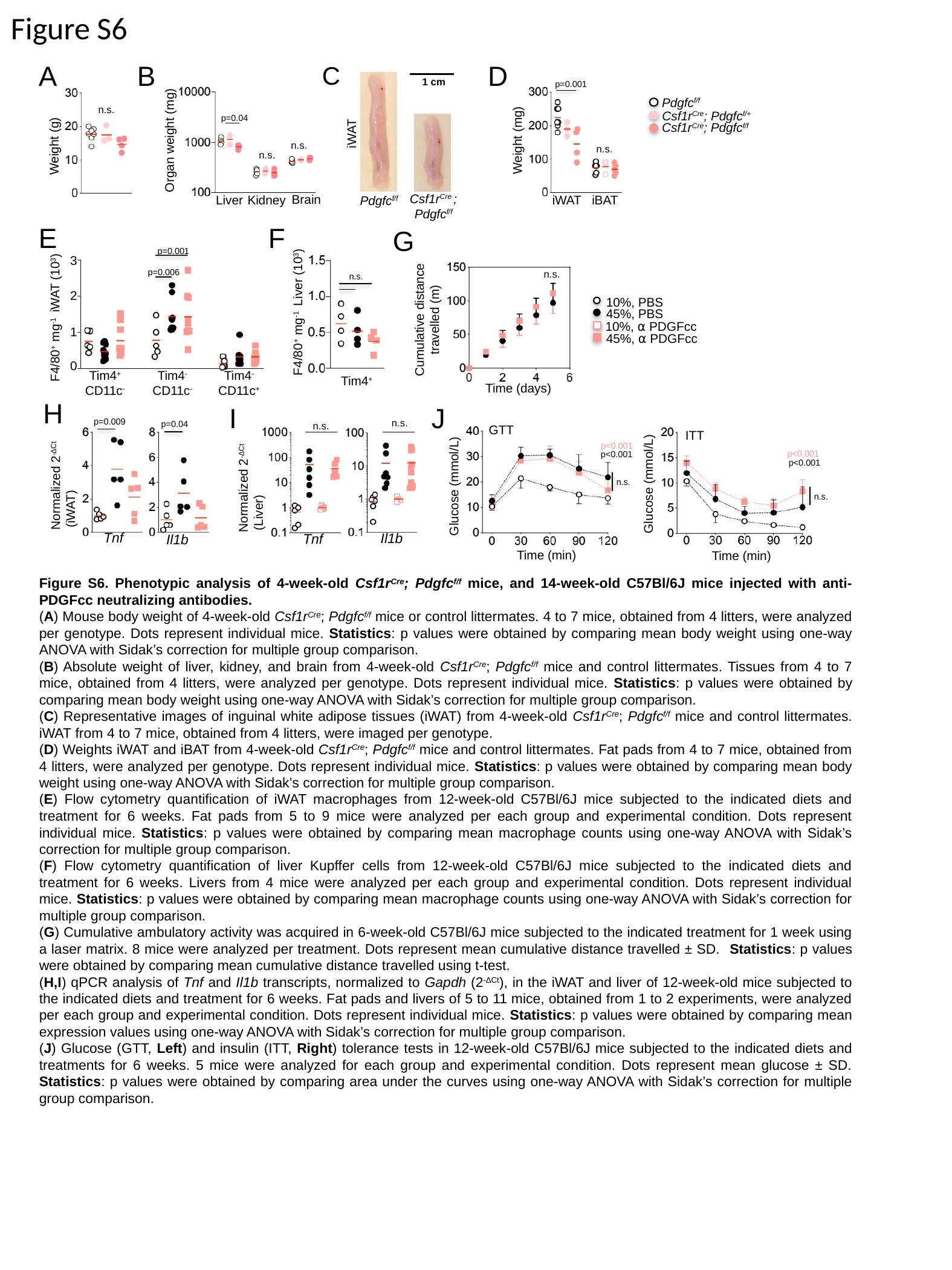

Figure S6
A
B
D
C
p=0.04
Organ weight (mg)
Brain
Liver
Kidney
1 cm
iWAT
Csf1r^Cre^ ; Pdgfc^f/f^
Pdgfc^f/f^
p=0.001
Weight (mg)
iBAT
iWAT
Pdgfc^f/f^
Csf1r^Cre^; Pdgfc^f/+^
Csf1r^Cre^; Pdgfc^f/f^
n.s.
Weight (g)
n.s.
n.s.
n.s.
E
F
G
n.s.
F4/80^+^ mg^-1^ Liver (10^3^)
Tim4^+^
p=0.001
3
p=0.006
2
F4/80^+^ mg^-1^ iWAT (10^3^)
1
0
Tim4^+^
CD11c^-^
Tim4^-^ CD11c^-^
Tim4^-^ CD11c^+^
Cumulative distance travelled (m)
Time (days)
n.s.
10%, PBS
45%, PBS
10%, ⍺ PDGFcc
45%, ⍺ PDGFcc
H
I
J
p=0.009
n.s.
p=0.04
n.s.
GTT
Normalized 2^-ΔCt^
 (iWAT)
Tnf
Il1b
Il1b
Tnf
ITT
p<0.001
p<0.001
p<0.001
p<0.001
Normalized 2^-ΔCt^
 (Liver)
n.s.
Glucose (mmol/L)
Glucose (mmol/L)
n.s.
Time (min)
Time (min)
Figure S6. Phenotypic analysis of 4-week-old Csf1r^Cre^; Pdgfc^f/f^ mice, and 14-week-old C57Bl/6J mice injected with anti-PDGFcc neutralizing antibodies.
(A) Mouse body weight of 4-week-old Csf1r^Cre^; Pdgfc^f/f^ mice or control littermates. 4 to 7 mice, obtained from 4 litters, were analyzed per genotype. Dots represent individual mice. Statistics: p values were obtained by comparing mean body weight using one-way ANOVA with Sidak’s correction for multiple group comparison.
(B) Absolute weight of liver, kidney, and brain from 4-week-old Csf1r^Cre^; Pdgfc^f/f^ mice and control littermates. Tissues from 4 to 7 mice, obtained from 4 litters, were analyzed per genotype. Dots represent individual mice. Statistics: p values were obtained by comparing mean body weight using one-way ANOVA with Sidak’s correction for multiple group comparison.
(C) Representative images of inguinal white adipose tissues (iWAT) from 4-week-old Csf1r^Cre^; Pdgfc^f/f^ mice and control littermates. iWAT from 4 to 7 mice, obtained from 4 litters, were imaged per genotype.
(D) Weights iWAT and iBAT from 4-week-old Csf1r^Cre^; Pdgfc^f/f^ mice and control littermates. Fat pads from 4 to 7 mice, obtained from 4 litters, were analyzed per genotype. Dots represent individual mice. Statistics: p values were obtained by comparing mean body weight using one-way ANOVA with Sidak’s correction for multiple group comparison.
(E) Flow cytometry quantification of iWAT macrophages from 12-week-old C57Bl/6J mice subjected to the indicated diets and treatment for 6 weeks. Fat pads from 5 to 9 mice were analyzed per each group and experimental condition. Dots represent individual mice. Statistics: p values were obtained by comparing mean macrophage counts using one-way ANOVA with Sidak’s correction for multiple group comparison.
(F) Flow cytometry quantification of liver Kupffer cells from 12-week-old C57Bl/6J mice subjected to the indicated diets and treatment for 6 weeks. Livers from 4 mice were analyzed per each group and experimental condition. Dots represent individual mice. Statistics: p values were obtained by comparing mean macrophage counts using one-way ANOVA with Sidak’s correction for multiple group comparison.
(G) Cumulative ambulatory activity was acquired in 6-week-old C57Bl/6J mice subjected to the indicated treatment for 1 week using a laser matrix. 8 mice were analyzed per treatment. Dots represent mean cumulative distance travelled ± SD. Statistics: p values were obtained by comparing mean cumulative distance travelled using t-test.
(H,I) qPCR analysis of Tnf and Il1b transcripts, normalized to Gapdh (2^-ΔCt^), in the iWAT and liver of 12-week-old mice subjected to the indicated diets and treatment for 6 weeks. Fat pads and livers of 5 to 11 mice, obtained from 1 to 2 experiments, were analyzed per each group and experimental condition. Dots represent individual mice. Statistics: p values were obtained by comparing mean expression values using one-way ANOVA with Sidak’s correction for multiple group comparison.
(J) Glucose (GTT, Left) and insulin (ITT, Right) tolerance tests in 12-week-old C57Bl/6J mice subjected to the indicated diets and treatments for 6 weeks. 5 mice were analyzed for each group and experimental condition. Dots represent mean glucose ± SD. Statistics: p values were obtained by comparing area under the curves using one-way ANOVA with Sidak’s correction for multiple group comparison.

### Slide 10
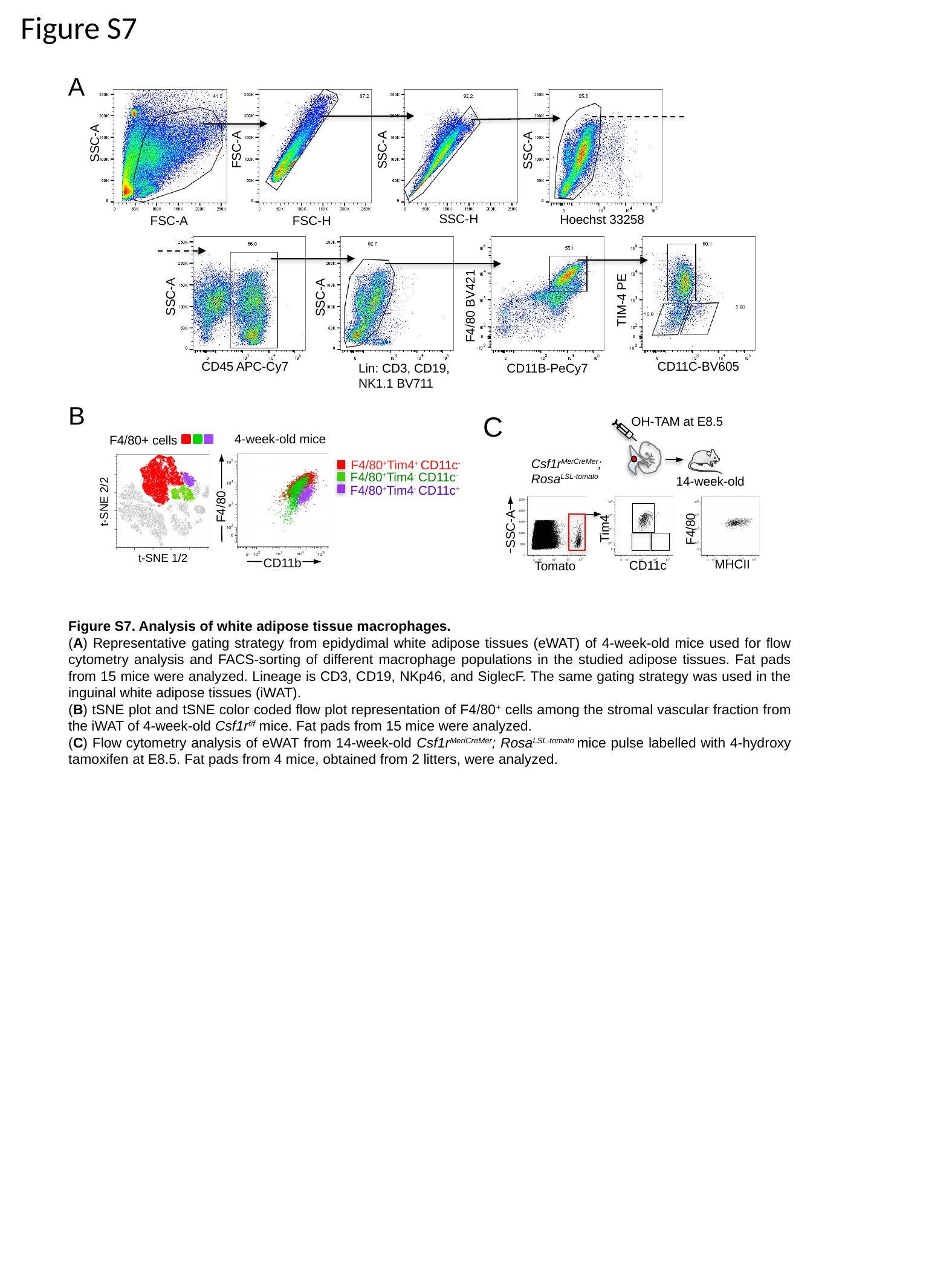

Figure S7
A
SSC-A
FSC-A
SSC-A
SSC-A
SSC-H
Hoechst 33258
FSC-A
FSC-H
SSC-A
SSC-A
TIM-4 PE
F4/80 BV421
CD45 APC-Cy7
CD11C-BV605
CD11B-PeCy7
Lin: CD3, CD19, NK1.1 BV711
B
C
OH-TAM at E8.5
Csf1r^MerCreMer^;
Rosa^LSL-tomato^
14-week-old
Tim4
CD11c
F4/80
SSC-A
MHCII
Tomato
4-week-old mice
F4/80+ cells
t-SNE 2/2
F4/80^+^Tim4^+^ CD11c^-^
F4/80^+^Tim4^-^ CD11c^-^
F4/80^+^Tim4^-^ CD11c^+^
F4/80
t-SNE 1/2
CD11b
Figure S7. Analysis of white adipose tissue macrophages.
(A) Representative gating strategy from epidydimal white adipose tissues (eWAT) of 4-week-old mice used for flow cytometry analysis and FACS-sorting of different macrophage populations in the studied adipose tissues. Fat pads from 15 mice were analyzed. Lineage is CD3, CD19, NKp46, and SiglecF. The same gating strategy was used in the inguinal white adipose tissues (iWAT).
(B) tSNE plot and tSNE color coded flow plot representation of F4/80^+^ cells among the stromal vascular fraction from the iWAT of 4-week-old Csf1r^f/f^ mice. Fat pads from 15 mice were analyzed.
(C) Flow cytometry analysis of eWAT from 14-week-old Csf1r^MeriCreMer^; Rosa^LSL-tomato^ mice pulse labelled with 4-hydroxy tamoxifen at E8.5. Fat pads from 4 mice, obtained from 2 litters, were analyzed.

### Slide 11
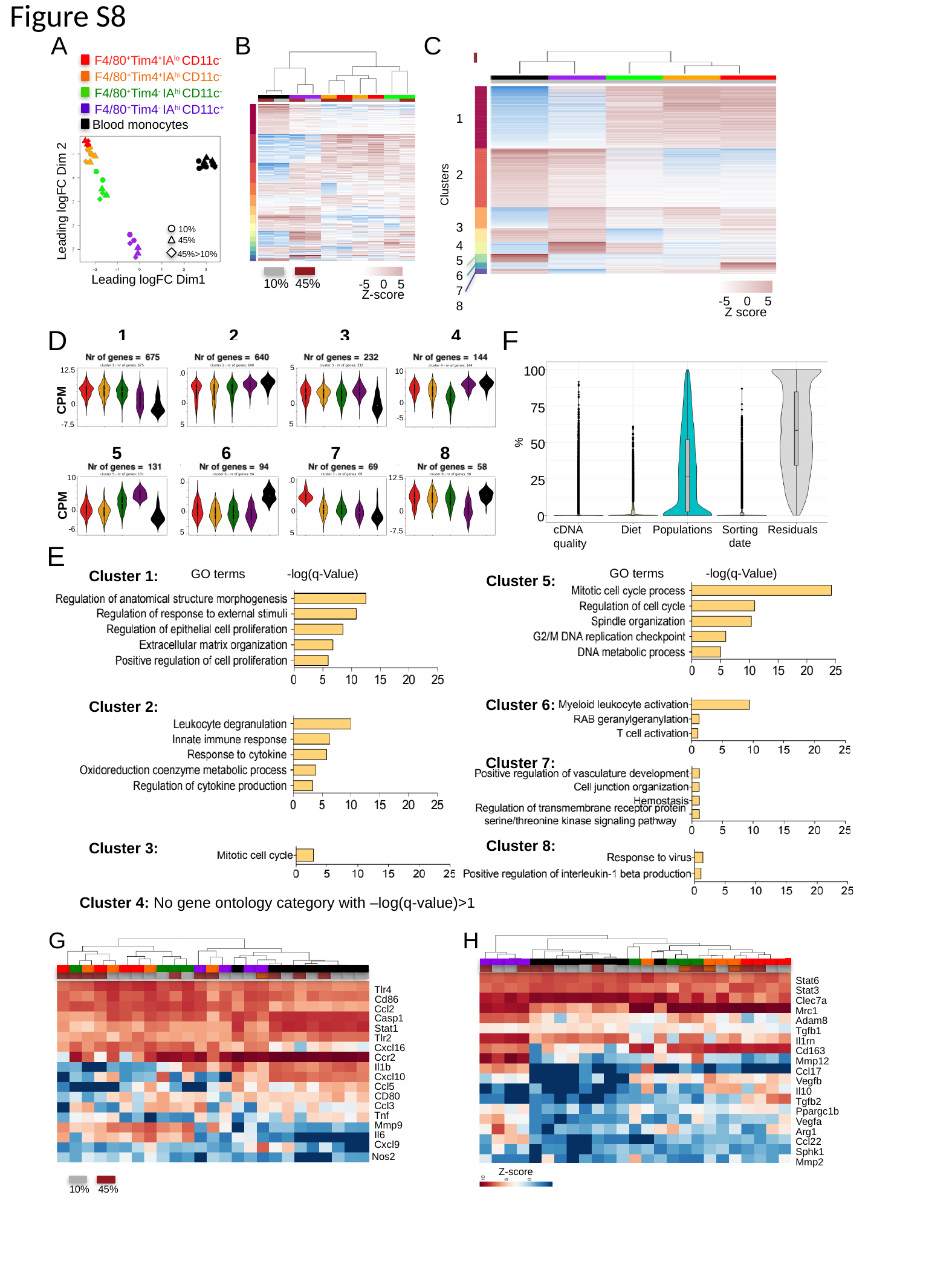

Figure S8
A
B
C
F4/80^+^Tim4^+^IA^lo^ CD11c^-^
F4/80^+^Tim4^+^IA^hi^ CD11c^-^
F4/80^+^Tim4^-^ IA^hi^ CD11c^-^
F4/80^+^Tim4^-^ IA^hi^ CD11c^+^
Blood monocytes
Leading logFC Dim 2
Leading logFC Dim1
10%
45%
45%>10%
1
2
3
4
5
6
7
8
Clusters
-5 0 5
Z score
10%
45%
-5 0 5
Z-score
1
2
3
4
12.5
12.5
10
10
CPM
0
0
0
0
-5
-7.5
-7.5
-7.5
5
6
7
8
10
12.5
10
10
CPM
0
0
0
0
-6
-7.5
-7.5
-7.5
F
D
100
75
50
25
0
Populations
Residuals
Diet
Sorting date
cDNA quality
%
E
GO terms
-log(q-Value)
GO terms
-log(q-Value)
Cluster 1:
Cluster 5:
45%
10%
Tim4^+^Ia^lo^ CD11c^-^
Tim4^+^Ia^hi^ CD11c^-^
Tim4^-^ Ia^hi^ CD11c^-^
Tim4^-^ Ia^hi^ CD11c^+^
Blood monocytes
Cluster 6:
Cluster 2:
Cluster 7:
Cluster 8:
Cluster 3:
Cluster 4: No gene ontology category with –log(q-value)>1
G
H
Stat6
Stat3
Clec7a
Mrc1
Adam8
Tgfb1
Il1rn
Cd163
Mmp12
Ccl17
Vegfb
Il10
Tgfb2
Ppargc1b
Vegfa
Arg1
Ccl22
Sphk1
Mmp2
Tlr4
Cd86
Ccl2
Casp1
Stat1
Tlr2
Cxcl16
Ccr2
Il1b
Cxcl10
Ccl5
CD80
Ccl3
Tnf
Mmp9
Il6
Cxcl9
Nos2
Z-score
10%
45%

### Slide 12
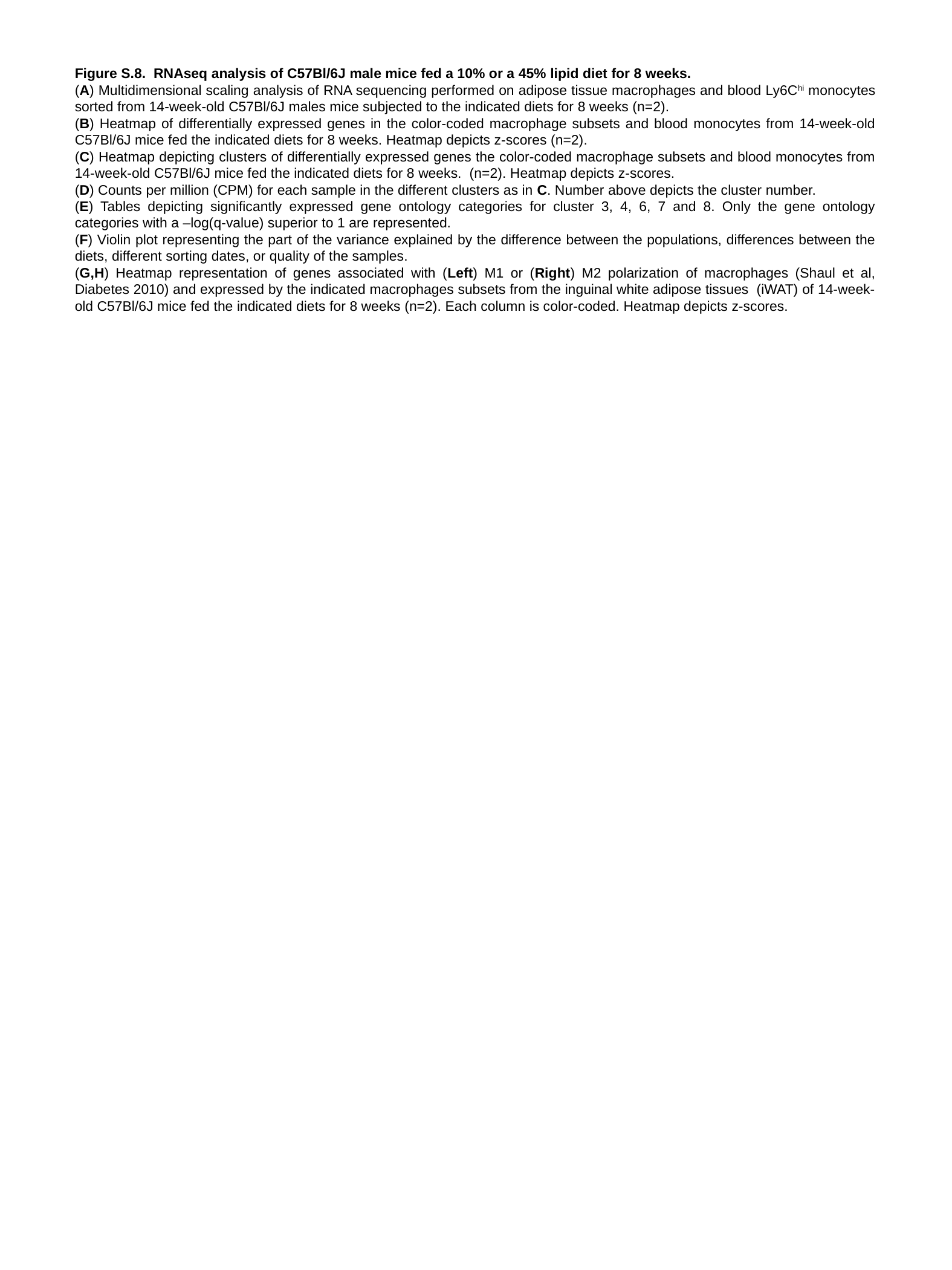

Figure S.8. RNAseq analysis of C57Bl/6J male mice fed a 10% or a 45% lipid diet for 8 weeks.
(A) Multidimensional scaling analysis of RNA sequencing performed on adipose tissue macrophages and blood Ly6C^hi^ monocytes sorted from 14-week-old C57Bl/6J males mice subjected to the indicated diets for 8 weeks (n=2).
(B) Heatmap of differentially expressed genes in the color-coded macrophage subsets and blood monocytes from 14-week-old C57Bl/6J mice fed the indicated diets for 8 weeks. Heatmap depicts z-scores (n=2).
(C) Heatmap depicting clusters of differentially expressed genes the color-coded macrophage subsets and blood monocytes from 14-week-old C57Bl/6J mice fed the indicated diets for 8 weeks. (n=2). Heatmap depicts z-scores.
(D) Counts per million (CPM) for each sample in the different clusters as in C. Number above depicts the cluster number.
(E) Tables depicting significantly expressed gene ontology categories for cluster 3, 4, 6, 7 and 8. Only the gene ontology categories with a –log(q-value) superior to 1 are represented.
(F) Violin plot representing the part of the variance explained by the difference between the populations, differences between the diets, different sorting dates, or quality of the samples.
(G,H) Heatmap representation of genes associated with (Left) M1 or (Right) M2 polarization of macrophages (Shaul et al, Diabetes 2010) and expressed by the indicated macrophages subsets from the inguinal white adipose tissues (iWAT) of 14-week-old C57Bl/6J mice fed the indicated diets for 8 weeks (n=2). Each column is color-coded. Heatmap depicts z-scores.

### Slide 13
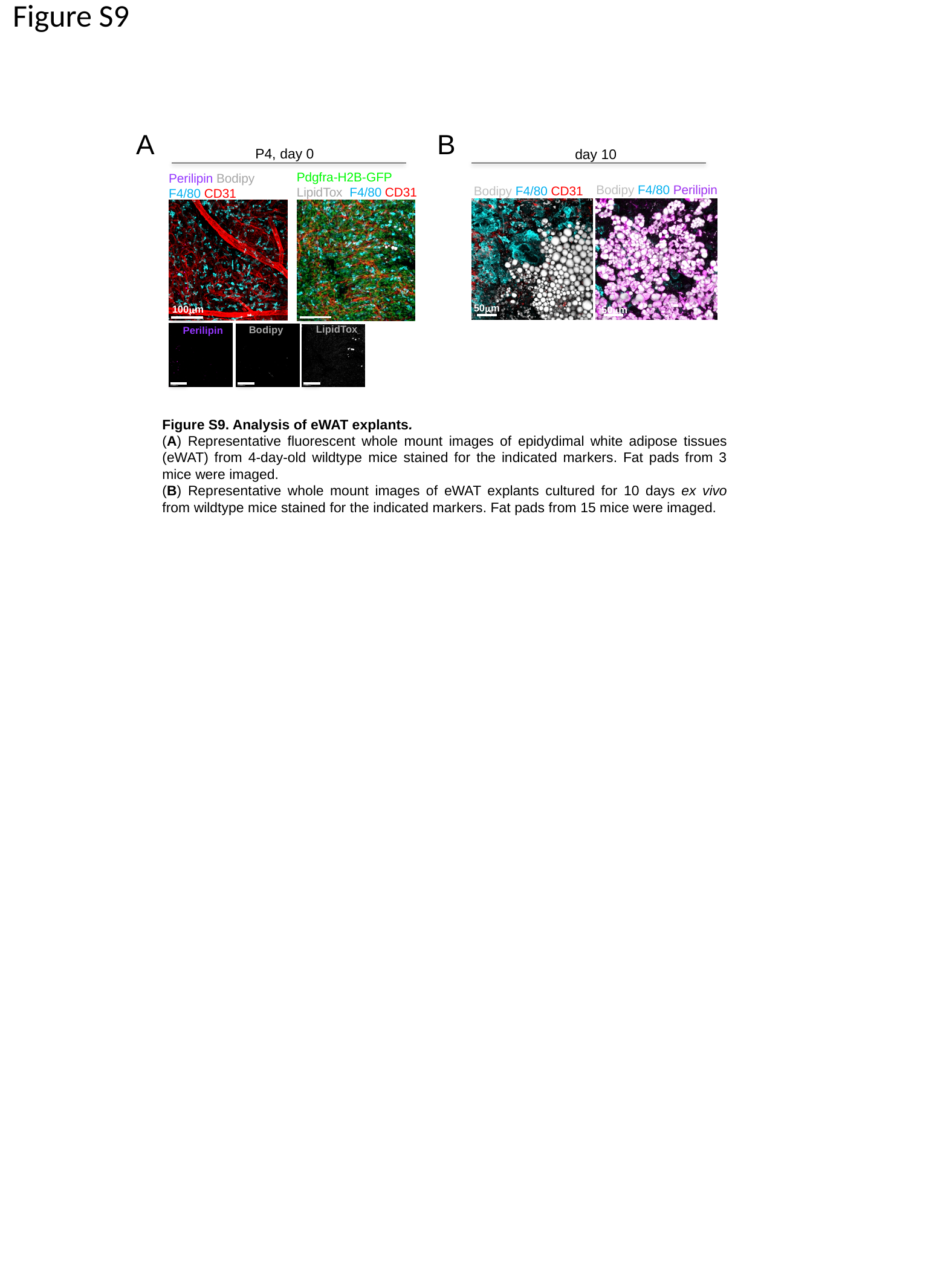

Figure S9
B
A
P4, day 0
day 10
Pdgfra-H2B-GFP
LipidTox F4/80 CD31
Perilipin Bodipy
F4/80 CD31
100μm
LipidTox
Bodipy
Perilipin
Bodipy F4/80 Perilipin
Bodipy F4/80 CD31
50μm
50μm
Figure S9. Analysis of eWAT explants.
(A) Representative fluorescent whole mount images of epidydimal white adipose tissues (eWAT) from 4-day-old wildtype mice stained for the indicated markers. Fat pads from 3 mice were imaged.
(B) Representative whole mount images of eWAT explants cultured for 10 days ex vivo from wildtype mice stained for the indicated markers. Fat pads from 15 mice were imaged.
